# PSTP: Decoding Latent Sequence Grammar for Protein Phase Separation through Transfer Learning and Attention

**DOI:** 10.1101/2024.10.29.620820

**Authors:** Mofan Feng, Liangjie Liu, Zhuo-Ning Xian, Xiaoxi Wei, Keyi Li, Wenqian Yan, Qing Lu, Yi Shi, Guang He

**Author notes:** **Corresponding author**: Qing Lu, Yi Shi, Guang He.

## Abstract

Phase separation (PS) is essential in various biological processes, necessitating high-accuracy predictive algorithms for studying numerous uncharacterized sequences, and accelerating experimental validation. However, many recent prediction methods face challenges in generalizability due to their reliance on engineered features. Furthermore, accurately identifying protein regions involved in PS remains challenging. To address this, we propose PSTP, a model employing a dual-language model embedding strategy and a lightweight attention module. The attention layer enables reliable residue-level phase separation predictions, identifying 84% of PS regions in PhaSePro and substantially improving correlation coefficient compared to existing models. PSTP also demonstrates robust performance in predicting PS propensity across various types of PS proteins and shows potential for predicting artificial proteins. By analyzing 160,000+ variants, PSTP characterizes the link between the incidence of pathogenic variants and residue-level PS propensities. PSTP’s predictive power and broad applicability make it a valuable tool for understanding biomolecular condensates and disease mechanisms.

## Introduction

Biomolecular condensates formed through phase separation (PS) are found throughout eukaryotic cells^1, 2, 3^, enabling a wide range of functions across multiple scales^4^. For example, at the molecular level, they regulate biochemical reaction rates such as photosynthesis^5, 6^ and mRNA degradation^7^, and participate in immune signaling^8, 9, 10^ and macromolecule folding^11, 12, 13^. At the mesoscale, condensates organize protein densities at neuronal synapses^14, 15^ and the nucleolus^16, 17^, and help coordinate DNA damage response^18^. At the cellular level, condensates function to mediate subcellular localization^19, 20, 21, 22^, and buffer against stochastic cellular noises^23, 24^. Dysregulation of PS is linked to numerous diseases, including cancer and neurodegeneration^25, 26^.

PS is a complex and not fully understood process^27, 28^, driven by multivalent interactions involving various physicochemical forces^29, 30, 31, 32, 33^. Both structured regions and intrinsically disordered regions (IDRs) contribute to driving PS^1, 2^. Recent studies on IDR ensembles have revealed a strong relationship between the conformational properties of IDRs, particularly their chain compaction, and their ability to drive PS^34, 35, 36^. Considering the vast amount of uncharacterized sequence data, developing reliable methods to predict PS and identifying the sequences responsible for this process is crucial for advancing our understanding of biomolecular condensates, their functional roles, and the mechanisms underlying disease development.

To predict PS, ‘first-generation’ algorithms were developed, focusing primarily on specific protein sequence characteristics^29, 37, 38, 39, 40, 41^. More advanced, integrated feature-based models^42, 43, 44, 45, 46^ have since been introduced, combining multiple sequence features with novel attributes such as immunofluorescence imaging, post-translational modifications (PTMs), protein-protein interactions (PPIs), and functional annotations. Recently, models have emerged that predict co-phase separation^47^ and differentiating PS proteins from those forming amyloids^48, 49^.

Nevertheless, many recent high-performing models rely heavily on engineered features from various sources, such as protein annotations and imaging data, which limits their generalizability to proteins, including variants and isoforms—that lack such detailed annotations. Moreover, current methods that provide residue-level PS scores often focus on one or a few specific sequence properties, making it challenging to accurately predict PS-driving regions due to the complex and diverse sequence patterns involved. These limitations hinder progress in uncovering key biological mechanisms and highlight the need for more flexible and broadly applicable models. To address these critical issues, we developed a model, termed PSTP (**P**hase **S**eparation’s **T**ransfer-learning **P**rediction) (Fig. 1). PSTP utilizes a dual-language model embedding strategy (Fig. 1A) to process and encode protein sequences. Specifically, we employ one large language model^50^ to extract latent biological features encoded in protein sequences; along with a modified molecular dynamics simulations-based language model^34^, encoding the sequence grammar underlying the physical and dynamic conformational properties of IDRs. This sequence-based feature embedding shows robust performance in predicting both self-assembly PS proteins and those whose PS behaviors are regulated by partner components, such as other proteins or nucleic acids (PS-Part proteins).

**Fig. 1.**
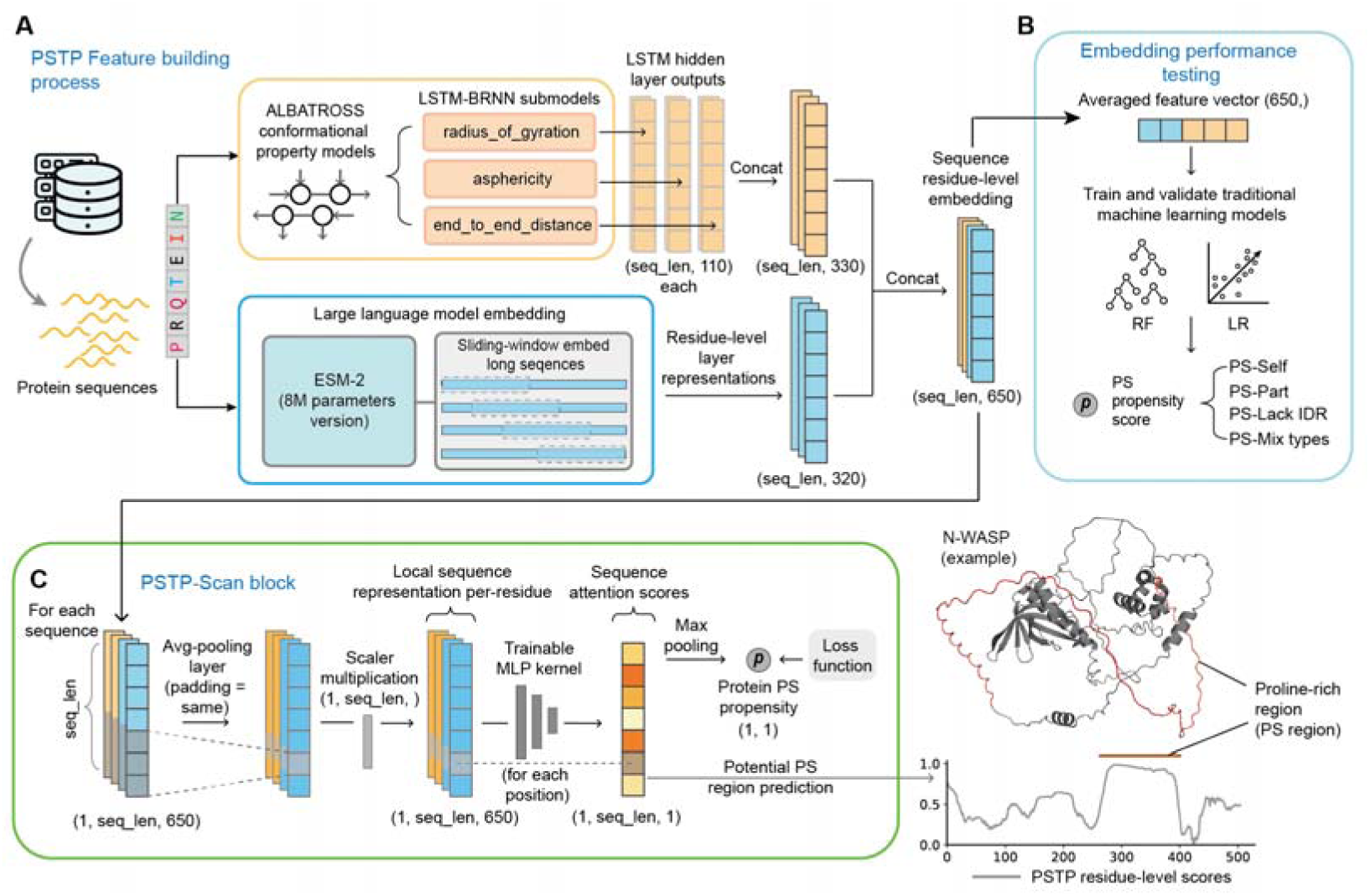
Schematic of PSTP embedding and machine learning (ML) architecture. (A) Protein sequences are converted into feature matrices using the hidden layer outputs of the ALBATROSS LSTM-BRNN and the final layer representations from the ESM-2 (8M parameters version) language model. For each sequence, the outputs from three ALBATROSS sub-models are extracted to form a matrix with dimensions of sequence-length x 330, while ESM-2 generates a matrix of sequence-length x 320. For sequences longer than 256, ESM-2 is applied in a sliding window manner to reduce time and memory costs. The combined result is a matrix of sequence-length x 650 for each sequence. sequence-length: length of the protein sequence. (B) Training and validation of traditional ML models using the PSTP embedding. The sequence-length x 650 embedding matrix is averaged to a vector of length 650 for each sequence. Four categories of phase separation (PS) proteins are considered: self-assembly PS proteins (PS-Self), partner-dependent PS proteins (PS-Part), mixed PS proteins (PS-Mix types), and PS proteins lacking IDRs (PS-Lack IDR). (C) Schematic of the PSTP-Scan block. The embedding matrix generated in (A) is used as input to produce attention scores, which serve as residue-level PS scores to detect PS regions. The maximum attention score for each sequence is output as the sequence-level PS propensity score. Here, we present N-WASP as an example, where PSTP-Scan predicts the proline-rich motif that forms multivalent interactions with its PS partner, Nck. MLP: multilayer perceptron.

Importantly, recognizing that PS is often driven by specific regions rather than the entire protein^51, 52, 53, 54, 55, 56^, we developed a lightweight attention-based model PSTP-Scan, to extract local information from sequence embeddings to focus on these key regions (Fig. 1C). Using 143 experimentally validated PS regions in PhaSePro^57^, we systematically evaluated existing residue-level PS predictors and found that PSTP-Scan’s attention weights significantly enhance the identification of PS-driving regions, without specific residue-level training. Notably, it identified 120 out of 143 PS regions, and improved the correlation coefficient to ∼150% of that of existing models.

In addition, we demonstrate the applicability and generalizability of PSTP across various protein types, such as artificial intrinsically disordered proteins (A-IDPs) and truncated proteins that undergo PS. Leveraging the high-throughput capability of PSTP, we characterized the relationship between pathogenicity and protein PS in low conservation regions, whose variants are much less studied compared to structured domains, using over 160,000 variants from ClinVar^58, 59^. We believe PSTP will facilitate a deeper understanding of cellular processes and aid in disease research and therapeutic development.

PSTP models were designed with ease of use and portability in mind. To ensure low memory consumption, we implemented a sliding window approach, making the large language model embedding and prediction within seconds on both standard CPUs and GPUs. We provide a locally installable version of PSTP, offering full access to the algorithms. Additionally, a user-friendly web server has been developed that enables prediction for easy prediction of sequences of interest.

## Results

### Phase separation proteins and IDR conformational properties

IDRs play important roles in driving protein phase separation^1, 2^ (PS), and recent findings^34, 35^ show that their ability to drive PS is closely tied to their degree of chain compaction. Specifically, compact IDRs favor PS^35, 60^, while expanded IDRs disfavor it^36, 61, 62^. Several sequence features, such as charge distribution^34, 63, 64^, hydrophobic^32,65^ patterning and aromatic residue content^32, 34, 35^, significantly influence the compaction of IDRs.

We first investigated different types of phase-separating proteins (PS proteins) and the conformational properties of their IDRs. We assessed the chain compaction of IDRs in various PS protein types by predicting their Flory scaling exponent (ν)^35^, the conformational entropy per residue (S_conf_/*N*)^35^, and Aspericity^34^ (Fig. 2 A and B), using models trained on molecular dynamics (MD) simulations data of IDRs. Higher values of these metrics indicate more expanded conformations, while lower values represent more compact ones. As expected, IDRs from PS proteins exhibited more compact conformations compared to those from the background proteome (Fig. 2 A and B). Notably, IDRs in proteins that self-assemble to form condensates (PS-Self proteins) are more compact than those in partner-dependent PS Proteins (PS-Part proteins), whose PS is regulated by partner components. This increased compaction correlates with the high PS propensity observed in PS-Self proteins. Besides IDRs, PS can also be driven by structured domains^2, 66, 67, 68^, and some proteins undergo PS in the absence of IDRs^69^. We collected 128 PS proteins that lack IDRs^69^ and only identified 28 IDRs from this group. These IDRs are significantly more expanded compared to other PS proteins, and even more so than those in the background proteome (Fig. 2 A and B). This expanded nature may facilitate multivalent interactions between structured regions that drive PS.

**Fig. 2.**
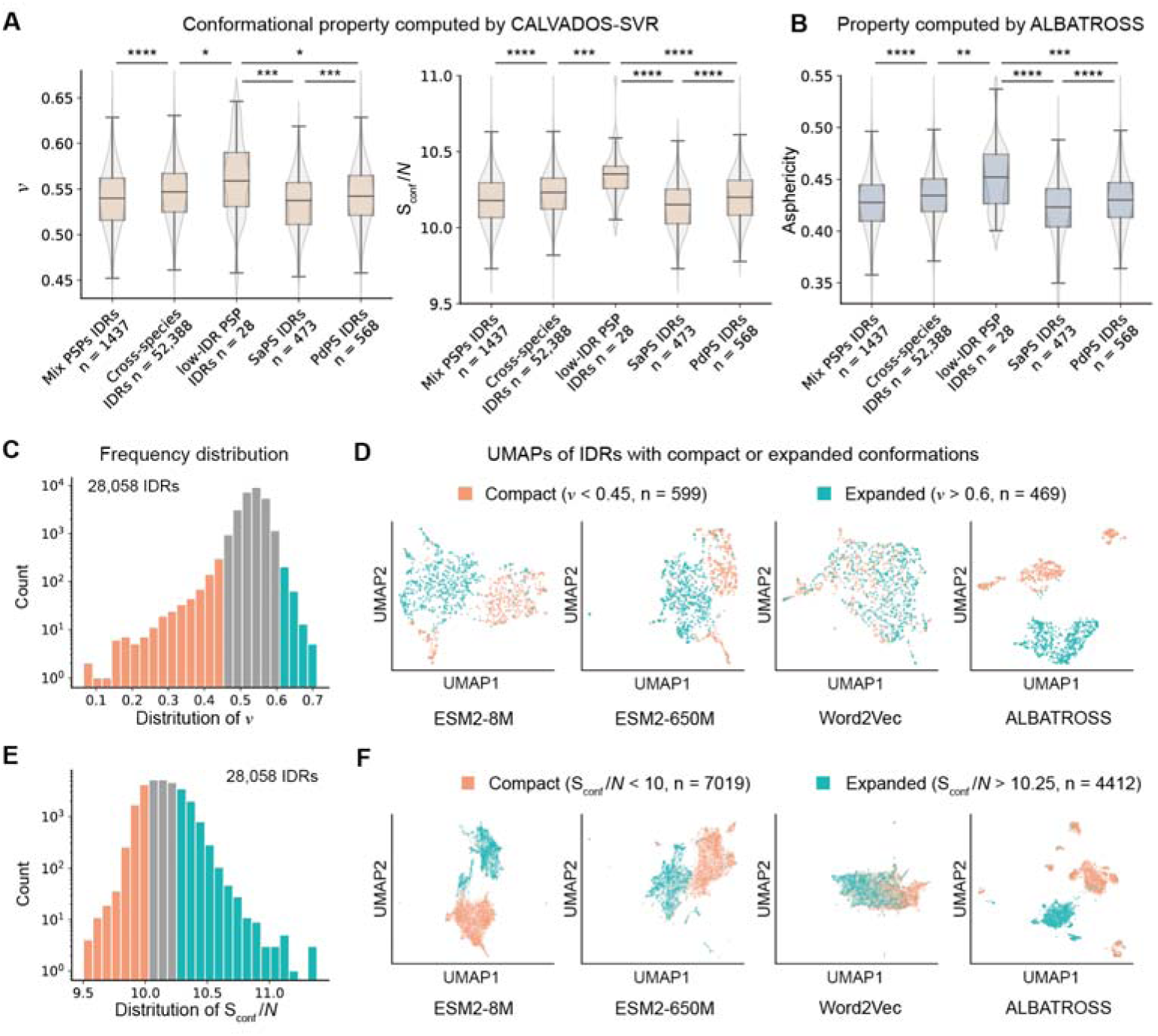
Analysis of IDR chain compaction properties and UMAP visualization of IDRs with varying chain compaction. (A and B) Comparison of conformational property metrics of intrinsically disordered regions (IDRs) across different protein types. Metrics reflecting IDR global conformation include *v* and S_conf_/*N* in (A), and asphericity in (B) (PSP: Phase separating proteins; ****P < 0.0001, ***P < 0.001, **P < 0.01, *P < 0.05; two-sided Mann–Whitney U test; boxplot components within each violin, from top to bottom are maxima, upper quartile, median, lower quartile, and minima.). (C) Distribution of ν values for IDRs. IDRs with ν < 0.45 are classified as extremely compact, while those with ν > 0.55 are classified as expanded. To balance data size, IDRs with ν > 0.6 were selected for the expanded group. (D) UMAP visualization of IDRs, categorized into compact (ν < 0.45) and expanded groups (ν > 0.6), illustrating the vectorized embeddings produced by different models. ESM2-8M and ESM2-650M represent the averaged layer representations from the 8M and 650M parameters versions of the ESM-2 model, respectively. Word2Vec represents embeddings produced by the skip-gram approach. ALBATROSS represents the averaged hidden layer outputs of the ALBATROSS model. (E and F) A parallel analysis to (C and D), considering IDRs with S_conf_/*N* < 10 as compact and IDRs with S_conf_/*N* > 10.25 as expanded.

### Protein embeddings capture IDR conformational property

In the previous section, we demonstrated the distinct conformational properties of IDRs across various types of PS proteins. Given that language model-based embeddings offer great flexibility and generalization in encoding protein sequences, and considering the relationship between IDR chain compaction and PS, we explored whether these models can effectively capture the conformational properties of IDRs. We collected the IDRome^35^ dataset, which contains 28,058 IDRs along with the conformation ensembles generated using CALVADOS^35, 36, 70^, an MD simulations model that shows a high correlation with experimental results. We then assessed whether language model-based embeddings could capture meaningful latent information related to sequence chain compaction by applying these methods to the IDRs. This was tested by visualizing the embedding representations for these sequences.

We first selected and colored extremely compact IDRs (ν < 0.45, n = 599) and expanded IDRs (ν > 0.60, n = 469) (Fig. 2C). Representations generated by the large language model ESM-2^50^ were able to distinguish between compact and expanded IDRs (Fig. 2D). Specifically, the visualization of the embedding space clearly shows that sequences are clustered into distinct groups, as demonstrated by both the smallest version of ESM-2 (8 million parameters) and the larger 650-million-parameters version. More pronounced distinctions were observed in the hidden output layer of ALBATROSS^34^, a deep neural network trained on MD simulation data (Fig. 2D, ‘ALBATROSS’).

We next evaluated the word2vec skip grams approach^71^ (Method), which was adopted in several PS prediction methods such as PSPredictor^72^, DeePhase^43^ and PSPHunter^44^. In contrast, we observe no significant differentiation between compact and expanded IDRs with this approach (Fig. 2D, ‘Word2vec’). We also visualized the vectors generated by an engineered-based approach based on van Mierlo et al.^42^, which considers amino acid composition, several dozen biophysical properties, and sequence low-complexity properties (a total of 52 features). We found that while compact and expanded IDRs are somewhat separated, the distinction is not significant as they still form overlapping clusters (Fig. S1). This shows the difficulties of engineered features for capturing the conformational information of IDRs.

We then divided the IDRs based on conformational entropy per residue (S_conf_/*N*), which is strongly correlated with ν. Using this metric, we classified IDRs as compact (S_conf_/*N* < 10, n = 7,019) or expanded (S_conf_/*N* > 10.25, n = 4,412) (Fig. 2E). The results were consistent with those observed using ν (Fig. 2F). These findings suggest that embeddings generated from both MD-based language models and the large language model capture meaningful information about IDRs, particularly with regard to chain conformational properties.

### Transfer learning contributes to phase separation prediction

Leveraging the conformational properties of IDRs and the underlying information decoded by protein large language models, we propose a residue-level protein feature embedding method (Fig. 1A). Specifically, for a given protein, we encode the IDRs’ chain conformation information by extracting the hidden layer outputs for three sub-ALBATROSS models trained on MD data to predict radius of gyration (Rg), end-to-end distance (Re), and asphericity, respectively. These outputs were concatenated to form a sequence-length x 330-dimension matrix (IDR conformation feature). In parallel, we incorporated the residue-level layer representations from the ESM-2 model (8-million-parameters version), resulting in a sequence-length x 320-dimensional matrix (large language model feature). To reduce memory and computational demands associated with long-distance attention calculations, a sliding window approach was applied for sequences longer than 256 residues (methods). By combining the outputs from ALBATROSS and ESM-2, we generated a sequence-length x 650-dimension feature matrix, referred to as PSTP (**P**hase **S**eparation’s **T**ransfer-learning **P**rediction) embeddings.

To assess PSTP embeddings’ effectiveness in encoding essential information for PS proteins, we first averaged the embedding matrix along the residue dimension (Fig. 1B). Using curated data from PhaSePred^45^, we trained and evaluated Logistic Regression (LR) and Random Forest (RF) models with default parameters to test the embeddings’ predictive power (Fig. 1B).

Cross-validation results show that PSTP embeddings achieved median AUCs of ∼0.89 and ∼0.80 with PSTP-LR for PS-Self and PS-Part, respectively, (Fig. 3A and B, S2A) demonstrating predictive ability across PS proteins from various species. To assess whether our dual-language embedding approach captures sufficient information, we tested its performance by adding both the word2vec embeddings and engineered features evaluated in the previous section. We observed that incorporating these features, or using them independently, did not improve performance, in some cases, even reduce it, as shown through cross-validation and independent testing with both LR (Fig. 3A) and RF (Fig. S2A). This shows the capability of PSTP embedding.

**Fig. 3.**
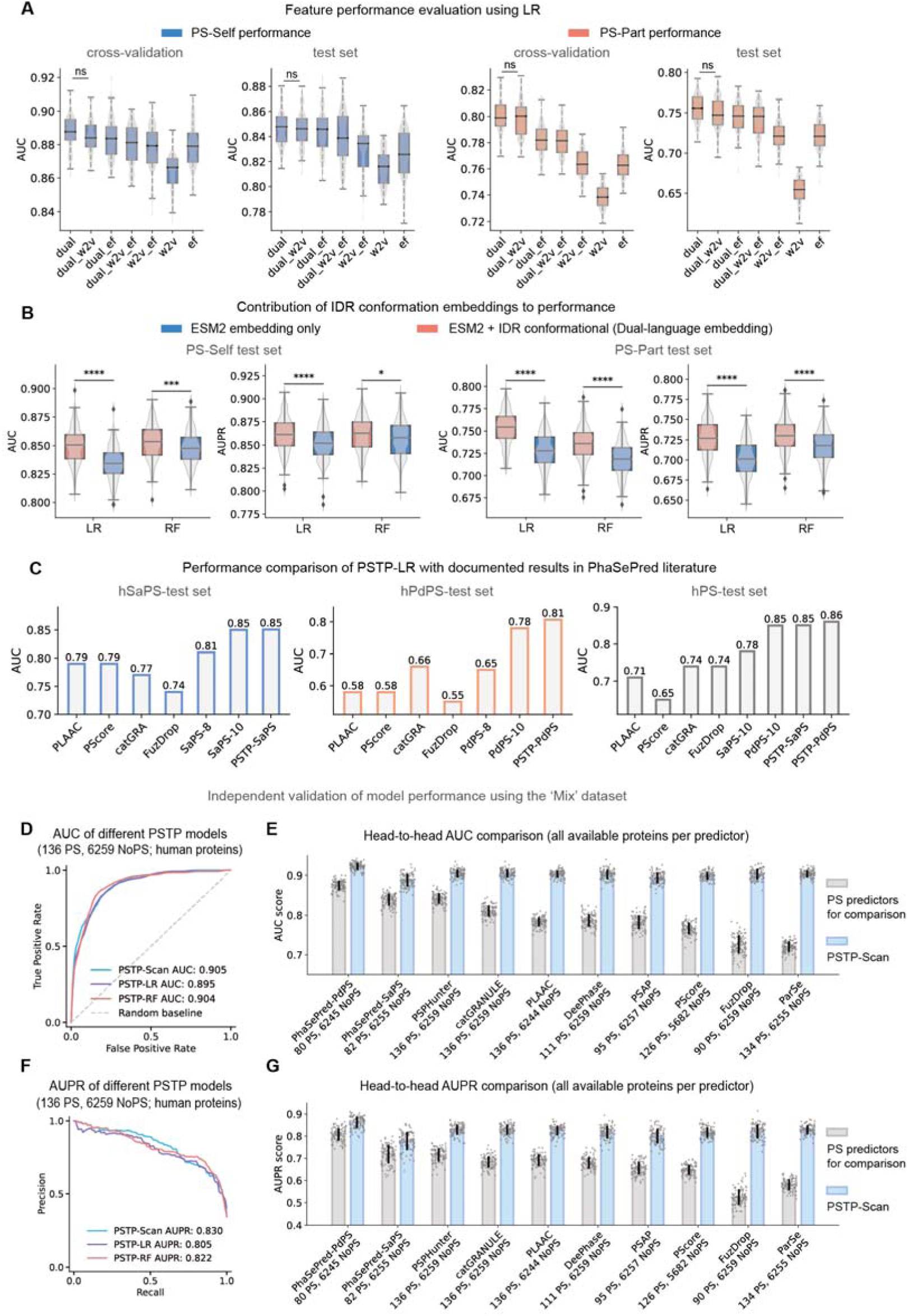
Evaluation of PSTP model performance. (A and B) Model performance of different feature combinations in predicting PS-Self and PS-Part proteins, evaluated through cross-validation and independent testing. Performance metrics were calculated using 50 replicates of subset sampling from the background dataset. (LR: Logistic Regression, RF: Random Forest; ****P < 0.0001, ***P < 0.001, *P < 0.05; two-tailed Student’s t-test; boxplot components within each violin, from top to bottom are maxima, upper quartile, median, lower quartile, and minima.). (A) Performance of different feature combinations using LR. ‘dual’: default PSTP-embedding, ‘w2v’: word2vec embedding, ‘ef’: engineered features. (B) Comparison of the default PSTP embeddings versus excluding IDR embeddings. Excluding IDR conformational embeddings (using only ESM-2 embeddings) reduced performance, while adding other features did not improve performance. (C) Comparison of PSTP-LR with PhaSePred (SaPS-8, SaPS-10, PdPS-8 and PdPS-10) as well as other representative models. PSTP-LR was trained and evaluated on the independent validation set following the same dataset split for PhaSePred. Results for PhaSePred and other PS predictors were sourced from the PhaSePred literature. (D) Model performances of different PSTP models on the independent validation dataset of PS-Mix proteins. Performance evaluated using 100 replicates of subset sampling from the background dataset is shown. (E) Head-to-head comparison between PSTP-Scan and other representative PS predictors. PSTP-Scan returns score for all proteins; the comparison with each method is thus performed on a set of proteins that can be predicted by that method, excluding those used in training for that method. AUC (Area under the Receiver Operating Characteristic curve) values were computed and visualized. Data are shown as mean ± SD (Standard Deviation), with scatter points representing performance across sampling repeats. (F and G) Parallel evaluations to (D and E) but focusing on the AUPR (Area Under the Precision-Recall Curve) values.

Notably, using only the IDR conformational feature (330-dimension vector) yielded only an average AUC of 0.81 for predicting PS-Self proteins and 0.72 for PS-Part proteins (Fig. S2C). Excluding IDR conformation features resulted in a significant performance drop across both cross-validation and the test set, underscoring the critical role of IDR conformational features in predicting PS (Fig. 3B).

Next, we independently validated PSTP-LR using the same training and independent validation set as PhaSePred, a state-of-the-art PS predictor. The SaPS-10 and PdPS-10 version of PhaSePred are meta-predictors that integrate multiple sequence-based algorithms, and multidimensional features such as immunofluorescence imaging and PTMs. Without additional feature selection or hyperparameter tuning, and using the sequence as the sole input, PSTP-LR achieved predictive accuracy comparable to SaPS-10 for PS-Self proteins, while showing an improvement in AUC (a 3% increase to PdPS-10) for predicting human PS-Part proteins (Fig. 3C). This PSTP-LR model also outperformed PhaSePred-10 on the hPS-test set, whether trained with PS-Self or PS-Part data (Fig. 3C, right panel).

The sliding window embedding approach with a length threshold of 256 residues for long sequences, did not lead to significant performance reduction. Increasing this threshold to 1024 residues (the maximum length during ESM-2 training^50^) showed no markedly accuracy gains (Fig. S2D), confirming the robustness of our initial configuration for handling longer sequences.

We further assessed PSTP’s performance in predicting membrane-less organelles (MLOs) participants localized to various intracellular locations using known MLO-associated protein datasets. These participants likely undergo PS or co-phase separation with other proteins to form MLOs. PSTP trained to predict PS-Part proteins demonstrated strong performance in identifying these proteins (Fig. S2F).

The high predictive accuracy underscores the significance of conformational features in PS and demonstrates the capability of pre-trained language models to capture the latent sequence grammar essential for PS.

### Model architecture of PSTP-Scan

In the previous section, the residue dimension-averaged PSTP embedding vectors showed robust performance. Since PS is often driven or regulated by one or a few specific regions within a protein, we suggest that leveraging the averaged feature vector of these local regions, rather than the full protein, may capture important information about these PS-prone areas. Drawing inspiration from image-based spatial attention neural networks, which generate attention scores for each pixel using local contextual information^73, 74, 75, 76^, we developed a model architecture named PSTP-Scan. This architecture provides attention scores that reflect residue-level PS profiles while simultaneously predicting the overall PS propensity of the protein.

For each sequence, the PSTP-Scan model takes the sequence-length x 650-dimension PSTP feature matrix as input. The attention weight for each residue position is generated using a convolution-like operation. This involves processing a local window of features around each position through an average pooling layer, followed by a trainable shared-weights two-layer multilayer perceptron (MLP) kernel (Fig. 1C, Method). The sequence-length unnormalized vector serves as the residue-level attention score.

The maximum attention weight represents the PS propensity of the region most likely to undergo PS, serving as the predicted PS propensity for the protein. This value is also used to update the shared-weights MLP through backpropagation during training, while the full vector of attention weights provides residue-level PS predictions.

Given that the lengths of PS-driving regions vary from tens to hundreds of residues (Fig. 4A), during training, the model employs multiple window sizes for the MLP kernel to capture information at different scales, enhancing robustness (Methods, Fig. S3C).

**Fig. 4.**
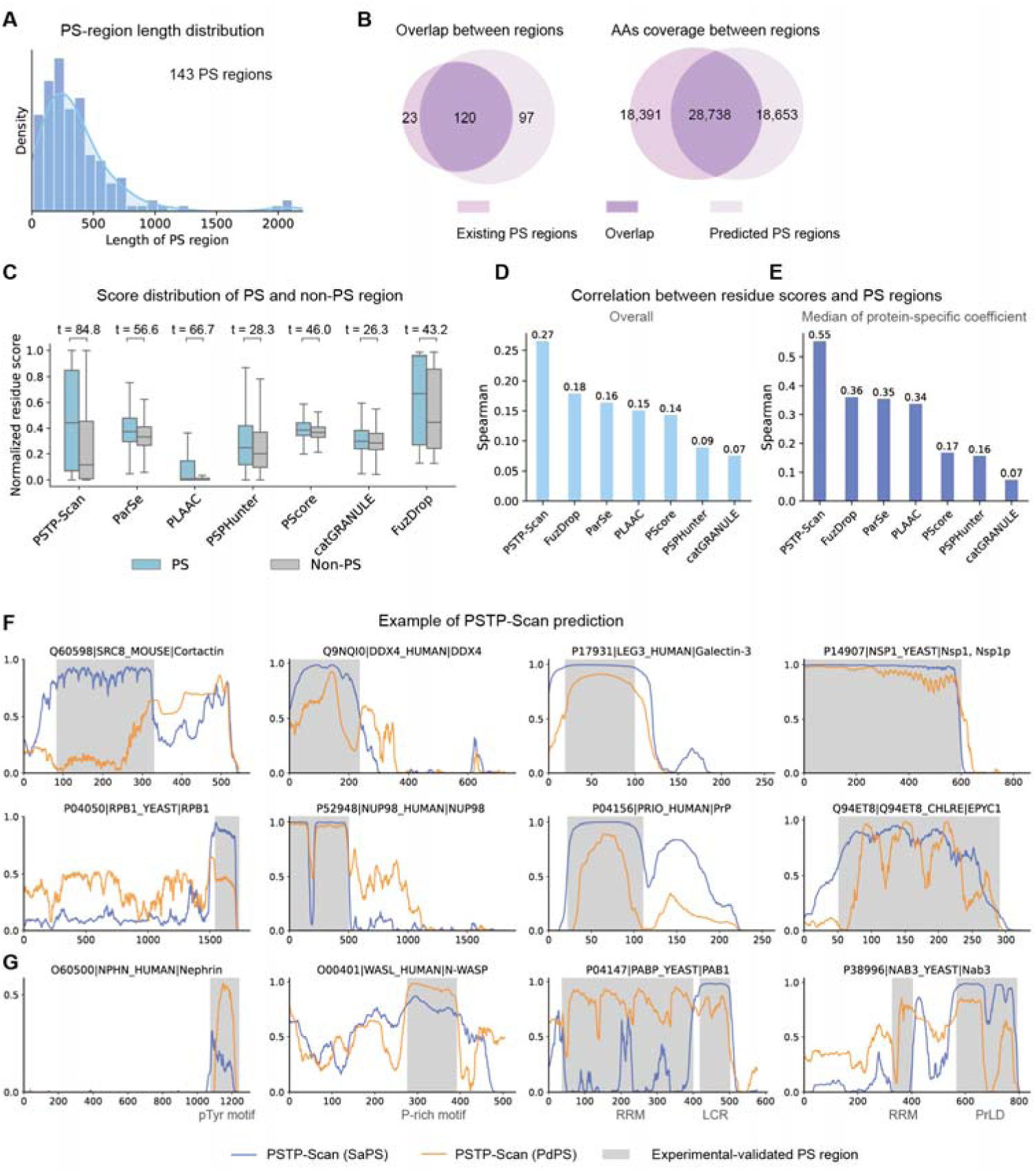
Evaluation of residue-level prediction scores generated by PSTP-Scan and other PS predictors under experimentally validated PS-driving regions in the PhaSePro database (143 PS regions from 121 proteins). (A) Distribution of sequence lengths for regions that drive PS. (B) Comparison of PS regions predicted by PSTP-Scan against those documented in PhaSePro. This panel highlights the overlap between predicted regions and the corresponding overlap at the individual amino acid level. (C) Distribution of the residue-level scores for PS regions versus non-PS regions in proteins documented in PhaSePro. All components have p-values < 0.0001. The boxplot components within each violin, from top to bottom are maxima, upper quartile, median, lower quartile, and minima. (D) Spearman correlation coefficient computed between residue-level scores and PS regions documented in PhaSePro for each method. The coefficients are computed between joint residue-level score vectors for each protein and a binary vector, where corresponding positions in PS regions are assigned 1 and others are assigned 0. (E) Median Spearman correlation coefficients at the protein level, computed between residue-level scores and PS regions for each protein. (F and G) Example of residue-level scores predicted by three PSTP-Scan models, with experimentally validated PS regions highlighted in gray. pTyr motif: phosphotyrosine motif; P-rich motif: Proline-rich motif; RRM: RNA-recognition motifs; LCD: Low-complexity domain; PrLD: Prion-like domain

### PSTP-Scan provides reliable PS prediction

Comparable to PSTP-LR, PSTP-Scan model also achieves robust performance in predicting both PS-Self and PS-Part proteins (Fig. S2E). This demonstrates that focusing solely on the most influential regions does not compromise the overall predictive accuracy.

To further evaluate PSTP’s robustness, we utilize a PS dataset curated by a more recent study^44^. The dataset (‘Mix’ dataset) consists of three PS sub-datasets, comprising a total of 892 PS proteins curated from various sources, including LLPSDB^77^, PhaSepDB^78^, DrLLPS^79^ and PhaSePro^57^, along with a background dataset of 8897 single-domain human proteins.

After applying CD-Hit^80^ to remove redundant sequences with a 0.4 threshold, we obtained a human-specific PS protein sub-dataset containing 136 proteins and a background set of 6259 human proteins (‘Mix’ test set), which served as the independent test set. We also obtained 375 PS proteins and 17,288 background proteins from other species, which were utilized as the training dataset (‘Mix’ training set, Methods).

PSTP-LR, PSTP-RF and PSTP-Scan all demonstrated strong performance in distinguishing human PS proteins from the human background proteins (Fig. 3D and F, Fig. S2G). We also evaluated individual the performance of a single PSTP-Scan block with different window sizes and observed consistently robust results (Fig. S2H), showing the robustness of PSTP-Scan, and insensitivity to window size variation.

We then conducted a head-to-head comparison of PSTP-Scan with current representative PS methods, excluding proteins used in the training datasets of those methods from the evaluation. The methods compared included PhaSePred^45^, which utilizes multi-dimensional integrated features; DeePhase^43^, which combines a word2vec language model with engineered features; PLAAC^37, 38^, which predicts prion-like propensities; PSAP^42^, which utilizes engineered biochemical and biophysical features; and ParSe^41^, a structure-based PS predictor. For comparison with PSPHunter^44^, we constructed features following its sequence encoding method. Additionally, we evaluated other sequence-based predictors, including catGRANULE^39^, FuzDrop^40^ and PScore^29^. PSTP-Scan achieves an AUC of ∼0.9 for all comparisons, better than these representative methods (Fig. 3 E and G).

These findings demonstrate that by leveraging large language models and deep learning models that encode IDR conformation information, PS-related features can be efficiently embedded. This approach provides a reliable and simplified method for PS prediction using only protein sequences as input.

The training and testing datasets in this section were combined together to form a ‘Mix’ dataset, that we used to train PSTP (Mix) model. We also trained a PSTP (SaPS) model using the PS-Self dataset as well as a PSTP (PdPS) model using the PS-Part dataset.

### PSTP-Scan improves identification of PS driving region

To evaluate PSTP-Scan’s ability to predict PS-driving regions at the residue level, we assessed the residue-level PS scores (the attention layer output) generated by PSTP-Scan (Fig. 1C). We utilize the experimentally validated PS regions documented in the PhaSePro^57^ database, derived from proteins across different species. Although PSTP-Scan’s training did not involve any residue-level information about PS regions, we specifically trained each sub-model evaluated here (SaPS, PdPS and Mix) after excluding proteins already included in the PhaSePro database (Fig. S4A) to avoid potential self-validation.

We found that, out of 143 experimentally validated PS regions documented in PhaSePro, 120 overlapped with the PSTP-Scan’s PS regions, which is a notable improvement over the 109 overlaps achieved by FuzDrop, a model directly trained on PhaSePro PS regions (Fig. 4B, Fig. S4 C and D). In terms of amino acid-level overlap between predicted PS and experimentally validated regions, PSTP-Scan recovered 28,378 residues, compared to 25,014 residues recovered by FuzDrop (Fig. 4B, Fig. S4 C and D). Furthermore, PSTP-Scan outperformed PSPHunter, a method recently reported to predict key regions for PS (Fig. S4 C and D).

Next, we systematically compared PSTP-Scan with other residue-level PS predictors in discerning PS regions. T-test statics showed that, compared to other predictors, PSTP-Scan has the strongest discriminative power in distinguishing residues located in PS driving regions from those in non-PS regions within PS proteins (Fig. 4C). Additionally, we computed the Spearman correlation between the residue-level scores of each predictor and the experimentally validated PS regions in PhaSePro (where a position is assigned 1 if it belongs to a PS region and 0 if not) (Fig. 4D). We calculated both the overall Spearman correlation and protein-specific correlations for each protein. PSTP-Scan achieved significantly higher correlations compared to other predictors, reaching approximately 150% of FuzDrop’s performance. In terms of protein-specific correlation, PSTP-Scan also demonstrated much higher performance over other methods (Fig. 4E and Fig. S4B).

In Fig. 4 F and G, and Fig. S5, we present examples of the PSTP-Scan prediction results. The Scan-SaPS model, trained on PS-Self proteins, identifies regions that drive PS, such as the low-complexity domain (LCD) repeats in cortactin^53^, and the N terminus IDR of Ddx4, which contains containing repeating net charge block and FG, RG blocks^81^. The Scan-PdPS model, trained on PS-Part proteins, outperforms Scan-SaPS in predicting PS-driving regions that rely on interactions with partners. This includes the phosphotyrosine motif of the Nephrin and the proline-rich domain of N-WASP (Fig. 4G) which form multivalent interactions with the SH2 and SH3 domains of NCK, respectively^55^. For proteins that contain both RNA-recognition motifs (RRMs) and LCDs, such as PAB1^52^ and Nab3^54^, Scan-PdPS identifies the RRMs, while Scan-SaPS focuses on the LCDs (Fig. 4G).

We next extracted records for PS-Self and PS-Part proteins, and found that PSTP-Scan achieves an overall correlation of 0.4 for PS-Self proteins (58 records, Fig. S4 E and H), second is PLAAC, a method to screen prion-like regions. All models showed significantly reduced performance for PS-Part proteins, with PSTP-Scan still performing the best (50 records, Fig. S4 F and I). Lower performance for PS-Part proteins may result from some protein-protein interaction sites are also denoted as PS-regions in the PhaSePro database, the prediction of which requires prior knowledge of specific protein interactions. For instance, PSTP-Scan did not predict the tri-RG motif in Buc that interacts with the regulator Tdrd6a^82^, but successfully predicted the prion-like domain responsible for Buc’s self-assembly^51^ (Fig. S5B), which is absent in the PhaSePro database. Similarly, while PSTP-Scan predicted the tandem repeats for the scaffold protein EPYC1, neither PSTP-Scan nor FuzDrop predicted the alpha-helices of Rubisco that bind to EPYC1^56^ (Fig. S5C). Incorporating models that focus on predicting protein-protein interactions may provide better insights into these types of PS mechanisms.

The limited performance of current methods may also stem from the highly flexible definitions of PS-driving segments or key regions. For example, PSTP-Scan achieves a 0.43 correlation with PS regions identified by NMR experiments (Fig. S4 G and J), significantly higher than the overall correlation. Nevertheless, given that PSTP-Scan predicts diverse PS-driving regions without being specifically trained on distinct PS regions, its ability to directly generate these predictions effectively demonstrates its validity, offering detailed insights into protein PS behavior.

### PSTP-Scan provides a broader range of prediction

Studying truncated proteins is essential for understanding protein function and pathogenicity, as NMD-escaping mutations that fail to trigger NMD often result in the expression of truncated proteins^83^. We evaluate PSTP-Scan as well as other sequence-based predictors on a test dataset of 93 experimentally truncated PS proteins and 242 non-PS proteins by collecting data from a recent study^84^. To avoid information leakage, each PSTP model evaluated here (PSTP-SaPS, PSTP-PdPS and PSTP-Mix) was specifically trained by excluding proteins corresponding to the protein fragment in the test dataset from the training dataset (Fig. S6A). We predicted these proteins using PSTP-Scan as well as PSTP-Scan’s MLP kernel, and evaluated their performance (Fig. 5B and Fig. S6B). We observed that compared with other representative sequence-based predictors, PSTP achieves the highest performance in both AUC and AUPR (Fig. 5B), while the MLP kernel from the PSTP-Scan (Mix) achieves the best performance at an AUC of 0.88 (Fig. 5B). This demonstrates PSTP-Scan’s robustness and effectiveness in predicting PS in truncated proteins.

**Fig. 5.**
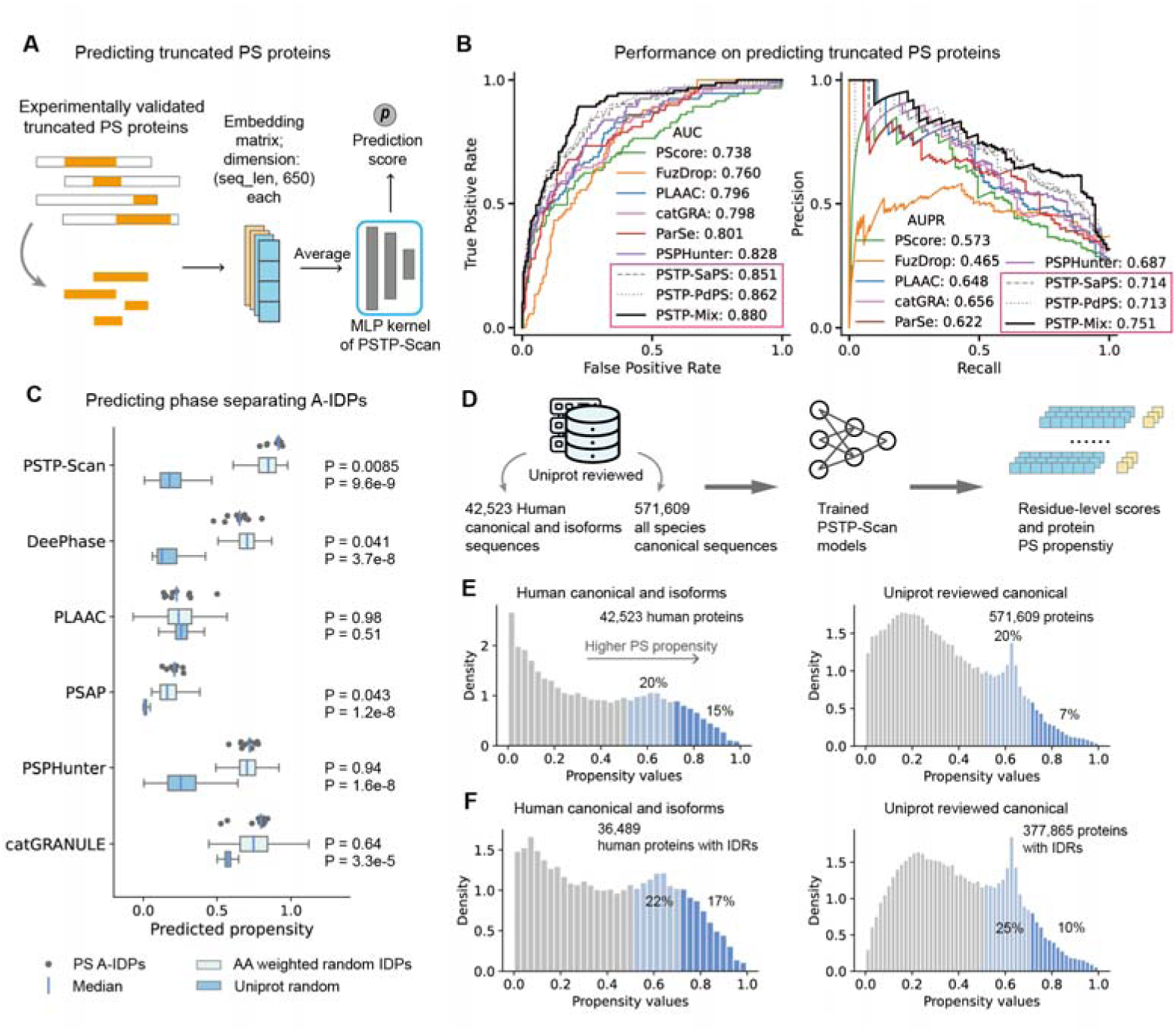
Extended scope of PS protein prediction. (A) Schematic of truncated protein prediction using the trained MLP kernel from PSTP-Scan. (B) Performance evaluation of different models for predicting experimentally validated truncated PS proteins and non-PS truncated proteins, assessed by AUC (left) and AUPR (right). (C) Score distribution comparison across models for predicting 11 artificial intrinsically disordered proteins (A-IDPs) that undergo PS, alongside random proteins. ‘Random’ represents scores for random 200-residue segments from UniProt-reviewed proteins (n = 10,000), while ‘AA weighted random’ refers to scores for control peptides, generated by repeating octapeptides 25 times as the A-IDPs, matching both the amino acid composition and repeating patterns of the A-IDPs. (D) PSTP-Scan-generated sequence-level and residue-level predictions for over 570,000 proteins from UniProt. (E) Distribution of PSTP-Scan scores across the human proteome, including both canonical proteins and isoforms (left), and for all UniProt-reviewed proteins (right). PSTP-Scan scores are determined by selecting the highest residue-level score from the three sub-models. (F) A parallel evaluation as (E), but with the inclusion of only proteins containing at least one IDR (IUPred > 0.5, length unrestricted).

In addition, a recent study^85^ designed artificial intrinsically disordered proteins (A-IDPs) to investigate the influence of amino-acid sequence patterns on the regulation of PS. This study created 11 artificial polypeptides based on prior knowledge from earlier research^86, 87^, each consisting of a 200-residue sequence formed by repeating an octapeptide 25 times, which exhibited PS at micromolar concentrations. Most of these A-IDPs underwent PS at room temperature, while some required higher temperatures.

The PSTP-Scan (Mix) kernel assigned high scores to these peptides. To compare, we generated 10,000 random background proteins by truncating 200-residue segments from UniProt-reviewed proteins. All models, except PLAAC, significantly distinguished A-IDPs from these background proteins, highlighting the unique sequence properties of the A-IDPs (Fig. 5C). However, PSAP produced relatively low scores, that failed to predict these peptides. Next, we generated control peptides by randomly repeating octapeptides 25 times, using the same amino acid composition as the A-IDPs (Fig. 5C, AA-weighted random). Interestingly, PSTP-Scan predicted lower scores for these control peptides (p-value = 0.0085), while other predictors did not show significant differences. This indicates that PSTP-Scan understands these A-IDPs by capturing information on both amino acid composition and the specific order of residues.

Both results indicate PSTP-Scan’s sensitivity to sequence information and its predictive power across a broad range of proteins. Leveraging the high-throughput nature of PSTP, we predict over 42,000 proteins in the human proteome, including canonical and isoforms, as well as over 570,000 Uniprot-reviewed proteins. We found that approximately 35% of human proteome and isoform proteins, and 27% of UniProt-reviewed proteins, contain potential PS regions (PSTP-Scan residue-level score > 0.5) (Fig. 5E). This proportion increases to 39% and 35%, respectively, when restricted to proteins with at least one IDR (Fig. 5F).

### Improved prediction of PSP that lack IDRs

To address the high disorder propensity (Fig. S7A) of current PS datasets, we also developed a predictor focused on PS scaffolds and regulators that lack IDRs. Utilizing a curated dataset from PSPire^69^, which includes 128 PS proteins and 10,251 background proteins, we trained and evaluated a PSTP-LR and a PSTP-RF model. Compared to PSPire, which utilizes structural information from AlphaFold2^88^, the PSTP models demonstrated greater robustness and reliability, achieving a ∼50% increase in AUPR (Fig. S7B and C).

### Incidence of pathogenic variants

Evolutionary conservation is currently a major factor in interpreting disease variants ^89, 90, 91, 92, 93^, and using this feature alone can predict the majority of known pathogenic variants^92^. IDRs tend to be less evolutionary conserved and lack fixed structures in comparison to structured and functionally essential regions^1^. As a result, studies on the pathogenicity of IDR variants have progressed slowly, leaving numerous variants of uncertain significance (VUSs) (∼ 530,000, ∼52% of ClinVar VUSs) unsolved in IDRs. Given the role of low-complexity IDRs in driving PS, we investigate the potential relationship between PS-driving propensity and the incidence of pathogenic variants in these low-evolutionary-conserved IDRs.

We analyzed PSTP-Scan residue-level scores for ClinVar variants located in low-conservation IDRs, defined by AlphaFold2 pLDDT score (pLDDT < 50) (Fig. 26A). The pLDDT scores, which reflects AlphaFold2 prediction confidence and is highly correlated with evolutionary conservation, is considered a state-of-the-art predictor for IDRs^35, 94^. We obtained 4756 pathogenic and 37,291 benign variants, and observed that, compared to benign variants, pathogenic variants tend to be located in positions with higher residue-level PS propensity (Fig. 6B). All sub-models of PSTP-Scan detected this difference, with Mann-Whitney p-values below 1 x 10^−10^. Examples of pathogenic IDR variants located in high-PS regions (above 0.8 in any PSTP-Scan sub-model) include P39L in HSPB1^95, 96^, F89I in DNAJB6^97^, and A315T in TARDBP^98, 99^, associated with Charcot-Marie-Tooth disease type 2 (CMT-2), limb-girdle muscular dystrophy type 1D (LGMD1D), and amyotrophic lateral sclerosis (ALS), respectively. These variants promote abnormal protein aggregation^95, 97, 98^ but are predicted as either VUS or benign by EVE^92^, an evolutionary-based top-performance pathogenicity predictor.

**Fig. 6.**
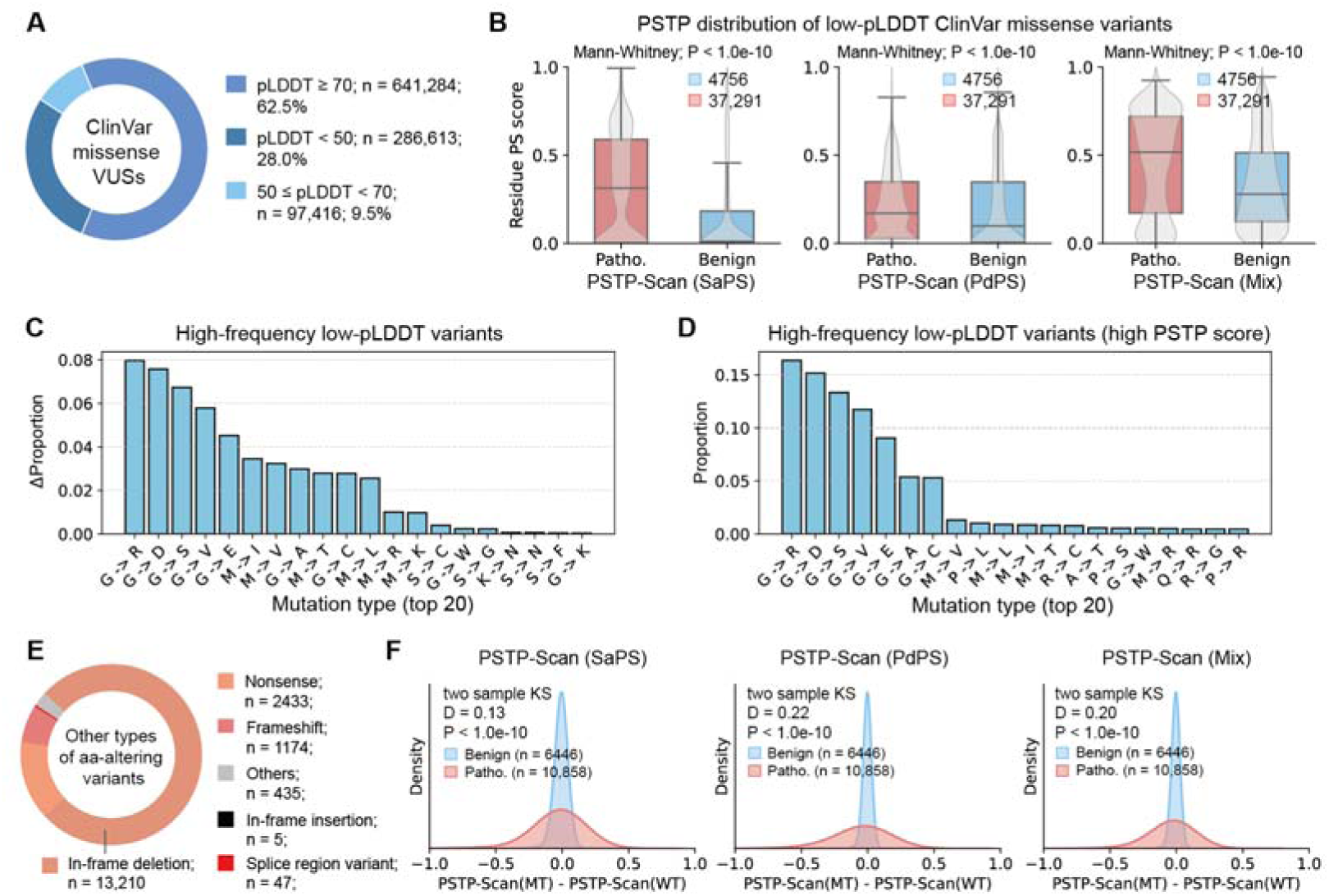
Analysis of the incidence of pathogenic variants and PS propensity. (A) Proportion of variants of uncertain significance (VUSs) based on pLDDT scores. (B) Distribution of PSTP-Scan residue-level scores for ClinVar variant positions with low-pLDDT (pLDDT < 50), classified as pathogenic (Pathogenic/Likely pathogenic) and benign (Benign/Likely benign). (C) Top 20 high-frequency mutations among low-pLDDT (pLDDT < 50) pathogenic variants. For each mutation type, ΔFrequency is computed as the difference in proportion between the low-pLDDT (pLDDT < 50) pathogenic group and other pathogenic variants (pLDDT ≥ 50). (D) Top 20 high-frequency mutations among low-pLDDT (pLDDT < 50) pathogenic variants with high PSTP-Scan residue-level scores (score > 0.5 from any sub-model). (E) Proportion of ClinVar protein-altering variants, excluding missense variants. (F) Distribution of PSTP-Scan score variations before and after mutations for pathogenic and benign ClinVar variants described in (E). Pathogenic variants show significantly higher PSTP-Scan score variations compared to benign variants.

A similar residue-level score distribution was observed for variants located in IDRs defined by IUPred3 (IUPred score > 0.5) (Fig. S6E). However, this distinction was weaker, possibly due to that IUPred-predicted IDRs also include evolutionary conserved IDRs^94^. Next, we evaluated this correlation based on allele frequency (AF), as variants with very low AF are likely to be pathogenic, while highly frequent variants are less likely to be^100, 101^. We collected variants from gnomAD V4.1.0 and found that, within low conservation regions, variants with low AF (AF < 1 x 10^−5^, 3,841,613 variants) tend to be located in positions with higher residue-level PS score region than variants with higher AF (AF ≥ 0.001, 25,673) (Fig. S6 D and F). In contrast, this correlation between PS and disease was not observed for variants located in structured regions (pLDDT ≥ 70) (Fig. S6C), where pathogenicity is influenced by factors including protein evolution and function.

We observed that within the disordered regions, particularly those with high phase separation (PS) propensity, glycine mutations are the most prevalent (Fig. 6 C and D), including neurodegeneration-related mutations such as G294V^102^ and G298S^103^ in TARDBP. These mutations, which replace Glycine with more rigid or charged residues, may alter the dynamics provided by Glycine and disrupt the original multivalent interaction pattern of specific key PS regions.

Our findings suggest that alterations in PS driving regions, are on average, more likely to lead to be pathogenic than alternation in non-PS regions, even though these PS driving regions are evolutionary less conserved.

In addition to missense variants, we also extracted 17,304 protein-altering variants, including nonsense, in-frame deletions, and frameshift variants (Fig. 6E). We computed the change in PSTP-Scan scores before and after mutation, and compared the distribution of these changes between pathogenic and benign variants using a two-sample KS test. We observed that pathogenic variants are more likely to alter PS propensity than benign variants, as indicated by the significant p-values of the KS test (Fig. 6F). This trend holds true both for the entire dataset (Fig. 6F) and when considering only the most frequent in-frame deletion mutation type (Fig. S6 G-I).

Our findings suggest that altering in PS driving regions, even in less-conserved disordered regions, are factors that contribute to pathogenicity. Combining effect on PS with traditional structure- and function-based pathogenicity analyses may enhance our understanding of disease mechanisms.

## Discussion

Protein language models have made significant progress in recent years. ESM-2 is a sequence-input-only encoder that provides a much faster and more flexible encoding compared to AlphaFold2^88^, which relies on computationally intensive multiple sequence alignments (MSAs). Although language-model-based embeddings may be less interpretable than engineered features, they capture comprehensive sequence characteristics more objectively, thus improving generalizability. Recent research shows the potential of the large language model in predicting PS^49, 104^. For broader applicability, we used the smallest version of the ESM-2 model and employed a sliding window approach to reduce memory usage, as its performance already demonstrates the effectiveness of the proposed approach.

While language models can extract rich evolutionary information from sequences, intrinsically disordered regions (IDRs) are relatively less conserved. Recent studies suggest that, while IDRs may undergo significant sequence variation throughout evolution, they can maintain stable conformational properties^34, 35^. Therefore, we consider that the conformational information of IDRs, which is closely linked to phase separation (PS) propensity, might be better captured by models specifically trained using MD simulation data. To address this, we designed an embedding approach that utilizes a pre-trained long short-term memory (LSTM) neural network to encode the dynamic properties of IDRs. While both types of embeddings alone can provide PS predictions using traditional machine learning models like logistic regression or random forest, their combination performs significantly better, confirming the effectiveness of our dual-language model embedding.

Unlike existing predictors such as FuzDrop^40^ or PScore^29^, which employ supervised learning models trained at each residue position, PSTP-Scan was trained without incorporating residue-level PS information, allowing it to learn autonomously. Due to the highly flexible definitions of PS-driving or regulating regions, the performance of current methods is often limited. However, PSTP-Scan’s design, which leverages local sequence information, makes it successfully predicts many PS driving segments, outperforming other models, and providing important insights into numerous uncharacterized sequences. PSTP-Scan also provides accurate protein-level PS prediction by focusing on the local region most likely to drive PS, aligning with the role of PS-driving regions in facilitating multivalent interactions with other protein regions^51, 52, 53, 82, 105, 106^.

Given that the training data size for phase-separating proteins is relatively small compared to other deep learning fields, to prevent overfitting, we avoided using overly complex deep models like transformer-based attention layers with query-key-value mechanisms^107^. We also chose not to use attention weights from the ESM-2 model, as they may emphasize structural and functional regions, introducing potential noise. Instead, we constructed an attention calculation method where only the weights of a multilayer perceptron (MLP) network are trainable.

To better handle dataset imbalance caused by using a large ‘background dataset’ as negative samples—a common training approach for current PS predictors^42, 43, 44, 45, 69^, we prevented the neural network from fitting the same data in every iteration. Specifically, in each epoch of PSTP-Scan training, we randomly select a small subset of proteins from the background dataset to serve as the negative dataset. This randomization helps prevent overfitting on background proteins that have not been experimentally confirmed as non-PS.

The disordered nature of IDRs not only makes the interpretation of their variants challenging but also limits the effectiveness of evolutionary or structure-based variant effect predictors like EVE^92^ or AlphaMissense^101^. This hinders the discovery and study of pathogenic variants. Our results reveal a link between the pathogenicity of low-conservation IDR variants and PS. Diseases related to PS often involve both abnormal protein aggregation and may also be linked to dosage effects^84^. To better interpret these variants’ pathogenicity, advanced methods are needed to capture and understand the evolution of IDRs, given their sequence divergence yet conformational conservation^34, 35^.

Biomolecular condensates perform essential cellular functions through complex and diverse interaction patterns. With its high performance, robustness, and sequence-only input nature, PSTP serves as a reliable tool for predicting biomolecular condensate formation. We believe PSTP will support a deeper understanding of cellular processes, aiding in disease research and therapeutic development.

## Methods

### IDR conformational property

#### Visualizing space distribution of IDRs of different properties

We collected curated datasets from several sources: self-assembly PS proteins (SaPS) (201 proteins), partner-dependent PS proteins (PdPS) (327 proteins) as well as non-redundant multi-species background proteins (60,220 NoPS proteins) from PhaSePred^45^. PS proteins that lack IDRs were collected from PSPire^69^ (128 proteins). In addition, we grouped PS proteins from several literatures^44, 45, 69^ to obtain a ‘Mix PSPs’ group (811 proteins after removing redundancy). For each group, we adopted IUPred3^108^ to predict residue-level disorder scores for each protein, following a process similar to Tesei et al.^35^ to filter IDRs. Specifically, residues with IUPred3 scores ≥ 0.5 were first labeled as disordered and those < 0.5 as folded. Regions shorter than ten residues were classified as gaps. These gaps were labeled as disordered if they were either flanked by disordered regions or terminal and adjacent to disordered regions. IDRs shorter than 30 residues or exceeding 1,500 residues were discarded.

The SVR model developed by Tesei et al.^35^ (https://github.com/KULL-Centre/_2023_Tesei_IDRome) is used to predict ν and S_conf_/*N* of IDRs. ALBATROSS developed by LotthammerLet al.^34^ (https://github.com/idptools/sparrow) is used to predict the asphericity of IDRs. The predicted conformational properties were compared between different IDR groups using the two-sided Mann–Whitney U test.

#### Visualizing space distribution of IDRs of different properties

The set of 28,058 IDRs used for visualization, along with their conformational properties generated by MD simulations^35^, were obtained from (https://sid.erda.dk/cgi-sid/ls.py?share_id=AVZAJvJnCO). Each IDR was vectorized using two models from ESM-2^50^: esm2_t6_8M_UR50D and esm2_t33_650M_UR50D, which output metrics with dimensions of sequence-length x 320, and sequence-length x 1280 respectively. Each matrix was then averaged into vectors of lengths 320 and 1280 for each IDR.

Additionally, a trained word2vec embedding method^109^ based on the skip-gram model was tested. The outputs for all contiguous 3-mer sequences of each IDR were summed to form a 100-length vector, which was also averaged to mitigate the impact of sequence length.

To evaluate the embeddings generated by ALBATROSS, we concatenated the hidden layer outputs from three sub-models for each input IDR into a sequence-length×330-dimensional matrix (see the ‘Feature Engineering’ section for more details) and averaged it to obtain a 330-dimensional vector.

Dimensionality reduction was performed using the UMAP algorithm, implemented through the Python ‘umap’ package.

### PS proteins dataset

#### Self-assembly PS and partner-dependent PS datasets

We utilize the curated datasets used for training and evaluating PhaSePred^45^, which were obtained from the supplementary materials provided with its published literature. A total of 201 SaPS proteins, 327 PdPS proteins, and 60,220 NoPS background proteins were collected. The acquired data included training and cross-validation (CV) datasets: SaPS (128 proteins), hSaPS (59 proteins), PdPS (214 proteins), hPdPS (96 proteins), NoPS (48,158 proteins), hNoPS (8801 proteins). The independent test sets comprised SaPS-test (73 proteins), hSaPS-test (34 proteins), PdPS-test (113 proteins), hPdPS-test (60 proteins), NoPS-test (12,062 proteins), hNoPS-test (2,200 proteins), PS-test (53 proteins), hPS-test (23 proteins).

#### Additional dataset for independent validation

To build an additional validation set, we utilize a PS dataset curated by a more recent study^44^. Among this data collection, we utilized the ‘hPS167’ datasets containing 167 human-specific PS proteins along with human background proteins as the independent test dataset, and the remaining PS proteins along with non-human background proteins as the training dataset. After reducing the sequence redundancy by CD-Hit at a threshold of 0.4, we obtained 136 human PS proteins, 375 non-human PS proteins and 6280 human background proteins. According to the ratio of non-human to human PS proteins, we randomly selected 2.75 times the size of the human background proteins from the NoPS dataset (60,220 proteins) proteins and obtained 17,270 proteins (PS proteins were filtered out). We next filtered both background proteins to ensure they were not longer than the longest human PS proteins (5,537 length), we finally got 6259 human background proteins and 17,268 non-human background proteins.

### Feature engineering

#### Feature embedding with protein large language model

We adopted two embedding matrices for each protein. The first matrix is the embedding computed by ESM-2^50^, we applied the esm2_t6_8M_UR50D (ESM2-8M in short) from the python ‘esm’ package: (https://github.com/facebookresearch/esm/tree/main.) ESM2-8M takes sequence as input and the sixth representation layer is output to return a 320-dimension vector for each position within each sequence. For each sequence, to reduce excessive memory usage as well as reduce the time required for long-distance attention computing, we apply a sliding window process similar to AlphaFold2^88^ and AlphaMissense^101^ when dealing with long sequences. Specifically, we applied a 256-length sliding window to scan through the sequence with a step size of 50 (For example, given a 520-length sequence, the sliding windows would be positioned at [1, 256], [51, 306], … [251,506], [265, 520]). For each residue position, if it appears in more than one window, we applied the residue embedding vector from the sliding window in which the position is most centered (farthest from both edges).

The speed of ESM2-8M is much faster compared to other ESM-2 versions on both CPU and GPU. Take an AMD Ryzen 7 3700X 8-Core Processor 3.7 G Hz CPU combined with 16 G memory as an example, for each 256-length-sequence (single sequence each batch), an average of 3.47 seconds is required for esm2_t6_8M_UR50D, while only an average of 0.11 seconds is required for esm2_t6_8M_UR50D (ESM2-8M). This makes it easier and faster for users to predict sequences using PSTP.

#### Feature embedding with ALBATROSS

The second embedding approach we adopted is the ALBATROSS program^34^ (https://github.com/idptools/sparrow). ALBATROSS is trained with MD simulation data of IDRs, and contains pre-trained LSTM-BRNN models to predict conformation properties. To capture as much conformational information as possible, we adopted three ALBATROSS submodels: the ‘asphericity’ model, the ‘radius_of_gyration_scaled’ model, and the ‘end_to_end_distance_scaled’ model. These three models share the same structure.

For each model of the three models, a protein sequence is inputted, and the hidden layer output of each timestep was adopted as the ALBATROSS embedding for each protein position, which returns a sequence-length x 110-dimension vector for each input sequence. These values range from −1 to 1. The vectors are then concatenated to a sequence-length x 330-dimension for each input sequence.

### Traditional machine-learning models

Two traditional machine learning models were applied to evaluate the feature embeddings we constructed. We applied a ‘LogisticRegression’ model using the ‘linear_model’ module and a ‘RandomForestClassifier’ model using the ‘ensemble’ module from the Python Sci-kit learn package. Both models were set with parameter class_weight = ‘balanced’ and other parameters were kept on default, and no normalization was applied to the dataset, neither was hyper parameter selection procedure. In each test scenario, to evaluate the LR and RF models during both 5-fold cross-validation (CV) and independent validation, the PS protein dataset was supplemented with randomly selected proteins (from the corresponding background dataset) at twice the size of the PS set. To ensure robustness, this sampling and validation process was repeated at least 50 times to calculate the average performance.

### PSTP-Scan neural network

#### Model architecture

PSTP-Scan takes the sequence-length x 650-dimension (320 for large language model embedding and 330 for IDR conformation embedding) feature matrix as input for each protein sequence. The model is implemented using the ‘pytorch’ package on Python.

**Within each PSTP-Scan block:** 1. The first layer is an average pool layer with ‘same’ padding, that for each protein position, a local window of feature matrix is averaged along the residue dimension to create a 650-dimensional vector. 2. To mitigate the effects of ‘same’ padding, a scaling step adjusts the values at the sequence ends to ensure they reflect the actual averages of the sequence elements rather than padded where the window size is 2*k* + 1, *x_j,d_* represents the *d*th value of the 650-dimensional input vector input for position *j*, and *L* represents the sequence-length:

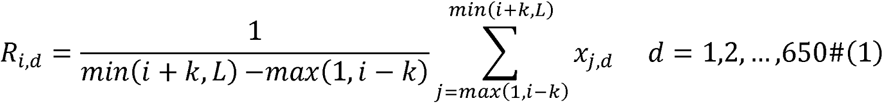

3. Next, a shared, trainable multilayer perceptron (MLP) kernel processes the averaged vector _i_. The MLP architecture consists of an initial layer with 20 neurons, followed by a LeakyReLU activation, a hidden layer with 5 neurons, and a final output layer with one neuron followed by a Sigmoid activation to produce a score between 0 and 1 for each residue position. These residue-level scores for each residue position form the sequence-length attention vector. So, the attention computation using the representation _i_ at each position can be written as the function (2). 4. The last is a max pooling layer that selects the maximum value of the attention vector, providing the sequence-level sequence propensity prediction, as shown in function (3).

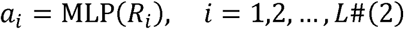

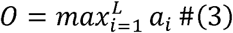

Given that the lengths of known phase-separating (PS) regions range from tens to hundreds of residues (Fig. 4A), three parallel blocks with identical architectures but different average pooling window sizes are used to capture PS information from sequence fragments of varying lengths. The pooling sizes are set to 256 + 1, 128 + 1, and 32 + 1, corresponding to powers of 2, with the extra 1 ensuring symmetric ‘same’ padding on both sides. We demonstrated the robustness of model performance across different window sizes (Fig. S2H) when a single PSTP-Scan block was applied for both training and validation. The predicted propensities (single-value sequence-level propensity) from these blocks are averaged to produce the final output, which is optimized through backpropagation to update the MLP parameters.

#### Model training

We initialized the network parameters using ‘kaiming_normal_’ with default settings. A modified LASSO regularization^110^ was applied to penalize the weights for ESM-2 embeddings (320 x dimension) and IDR embeddings (330 x *m*dimension) and in the first layer of the MLP, respectively, with the penalty 679 weight set *α* to 0.003 (shown as formula (4), where *W* represents the parameter weight matrix of the first

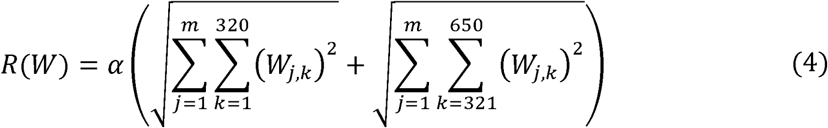

L1 regularization was applied to the other MLP layers, also with a penalty weight of 0.003. Binary cross-entropy loss was used. The Adam optimizer was applied with a learning rate of 0.003, without additional hyperparameter tuning. During the training process, for each iteration (50 epochs in total), we held out a random subset of proteins for the size of the positive dataset from the background dataset as the negative dataset. This approach was applied to address sample imbalance.

### Model performance benchmarks

Area Under the Receiver Operating Characteristic Curve (AUC), and Area Under the Precision-Recall Curve (AUPR) were applied during each validation. For the head-to-head comparison with PhaSePred and other PS predictors, we randomly selected proteins twice the size of the PS dataset from the background proteins and merged them with the positive dataset. Performance was evaluated using AUC and AUPR, and this process was repeated 50 times to calculate the average performance to ensure robustness.

### Evaluation of PS-Driving Region Identification

We collected PS proteins and their corresponding experimentally validated PS regions from the PhaSePro^57^ database (https://phasepro.elte.hu/download_full.tsv). To prevent self-validation, we trained each PSTP-Scan model (SaPS, PdPS and Mix) after excluding proteins already included in PhaSePro^57^. We set the average pool size at 33 (the minimum value of the average pool size when training PSTP-Scan) when generating the residue level scores for these proteins. We define the PSTP-Scan PS regions using the prediction values of the three models, that windows where scores were continuously over 0.5 more than 20 AAs by the maximum of three models were considered PS regions.

To systematically evaluate PSTP-Scan’s residue-level PS scores, we assigned residue-level scores to each protein based on the highest protein-level PS predicted value among the three sub-models: PSTP-Scan-Mix, PSTP-Scan-SaPS, and PSTP-Scan-PdPS. For example, if PSTP-Scan-PdPS yielded the highest predicted propensity for a protein—indicating it is likely to undergo partner-dependent PS—we used the residue-level scores from the PSTP-Scan-PdPS model for that protein.

We calculated Spearman correlation coefficients between these residue-level scores and the binary annotation of actual PS regions (assigning a value of 1 to positions within PS regions and 0 otherwise) to assess predictive performance.

### Final PSTP models

For the final PSTP (PSTP-LR, PSTP-RF and PSTP-Scan) models, we trained three versions, using the PS-Self data, PS-Part data and Mix PS data, respectively. The training and testing datasets for each task were combined to train the final model. For each of the three models, 10 sub-models were trained, and the average output of these 10 models was used as the final result at both the residue and protein levels.

### Other PS models

For each of PScore^29^, PLAAC^37^, PSAP^42^, ParSe^41^, and PSPHunter^44^, the prediction software was obtained from their respective GitHub repositories, as documented in the corresponding literature. PhaSePred^45^ scores for self-assembly PS proteins (‘SaPS-10fea_rnk’) and partner-dependent PS proteins (‘PdPS-10fea_rnk’) were downloaded from (http://predict.phasep.pro/download/). Prediction scores for FuzDrop^40^ were obtained from (https://fuzdrop.bio.unipd.it/). Prediction functions, as well as feature vectors and model parameters for catGRANULE^39^, were obtained from its corresponding supplementary information and methods. During the independent testing (Fig. 3 D to G), proteins that were included in the PS samples for each method were excluded from both positive and negative datasets.

### PS and Pathogenicity

For the pathogenicity analysis, we downloaded variants from ClinVar (https://ftp.ncbi.nlm.nih.gov/pub/clinvar/tab_delimited/variant_summary.txt.gz) as of April 2024, and from gnomAD V4.1.0 (https://gnomad.broadinstitute.org/).

### Data and code availability

All data mentioned in this paper is public data and can be obtained through the corresponding description in the results or methods. An installable Python package for the software is available at (https://github.com/Morvan98/PSTP).

## Acknowledgment

This project is supported by the National Key Research and Development Program (2022YFE0125300 to Y.S, 2020YFA0509700 to Q.L), Innovation Program of Shanghai Municipal Education Commission (2023ZKZD16) to G.H, the National Natural Science Foundation of China (82071262) to G.H, Shanghai Municipal Science and Technology Major Project (2017SHZDZX01 to Y.S, 20JC1418600 to G.H), Key Technology Breakthrough Program of Ningbo Sci-Tech Innovation YONGJIANG 2035 (2024Z221) to G.H, and Shanghai Jiao Tong University STAR Grant (YG2023ZD26, YG2023LC14, and YG2024QNA59 to G.H, YG2022ZD024, YG2022QN111 to Y.S, 23X010300421 to Q.L).

## Author contributions

M.F, Q.L, G.H and Y.S conceived the concept; M.F, L.L curated datasets, and designed computational algorithms and experiments; M.F, L.L, X.W, K.L and Q.Y performed computational experiments with assistance from Q.L, G.H and Y.S; M.F and Z.X analyzed data; G.H led the project with assistance from M.F, Q.L, Y.S, and X.W; M.F, Q.L, G.H and Y.S wrote the manuscript with input from all co-authors.

## Competing Interests Statement

The authors declare no competing interests.

## Supplementary figures

**Fig. S1.**
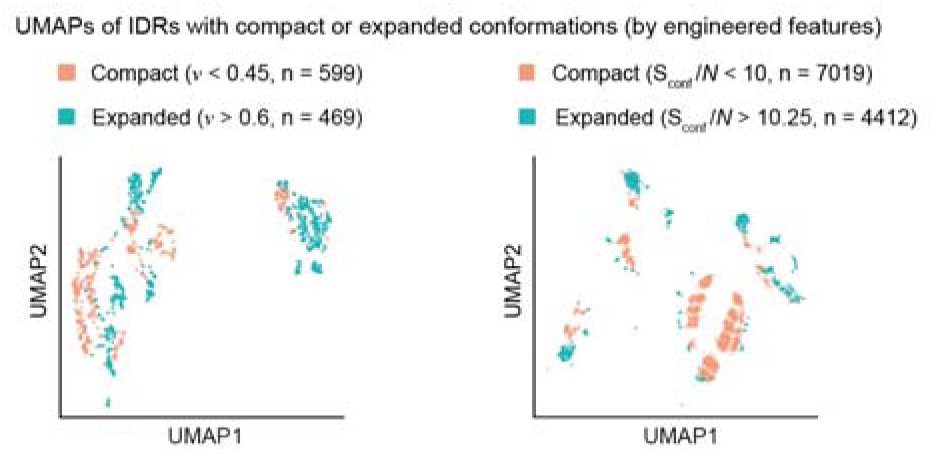
UMAP visualization of IDRs with varying chain compaction, related to Fig. 2. Parallel analysis to Fig. 2D (left) and 2F (right), with each IDR represented by engineered feature vectors incorporating various physicochemical properties.

**Fig. S2.**
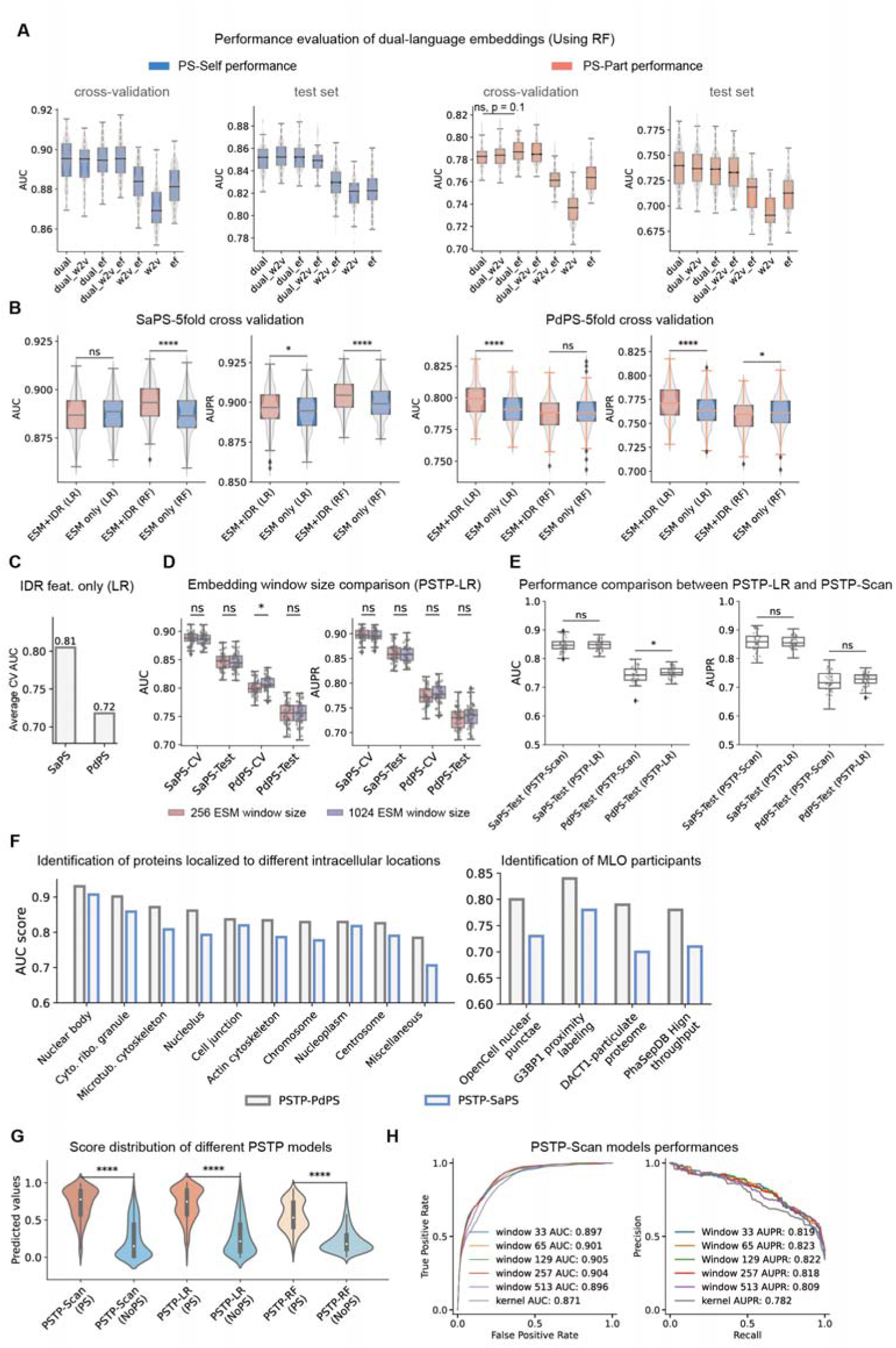
Model performance benchmark, related to Fig. 3. (A) Parallel evaluation to Fig. 3A, but using a random forest model to evaluate the performance of different feature combinations. ns: not significant. (B) Parallel evaluation to Fig. 3B, performed on the PS-Self (SaPS) and PS-Part (PdPS) cross-validation sets using 5-fold cross-validation. (C) Performance evaluation using only the 330-dimension IDR conformation features using the LR model under 5-fold cross-validation. (D) Comparison of sliding window embeddings (ESM-2 large language model embeddings) with 256-length and 1024-length thresholds across different cross-validation and independent testing. Scatter points represent the performance scores from repeated sampling. The performance of both window lengths is comparable. (*P < 0.05, ns: not significant; two-tailed Student’s t-test; boxplot components within each violin, from top to bottom are maxima, upper quartile, median, lower quartile, and minima.). (E) Performance comparison of PSTP-LR and PSTP-Scan on the SaPS and PdPS test set. Performance was evaluated using 50 replicates of subset sampling from the background dataset, with scatter points representing performance scores from these repeats. Both models show similar performance (ns: not significant, *P < 0.05; two-tailed Student’s t-test). (F) Prediction performance of PSTP-LR models trained on PS-Self data (PSTP-SaPS) and PS-Part data (PSTP-PdPS) in predicting MLO participants from those localized to intracellular locations (left) and four MLO datasets (right). These participants are curated from the PhaSePred literature. AUC values were calculated using proteins in these MLO datasets as positive samples and the human NoPS dataset as negative samples, excluding those in the model training data. (G) The discriminative power of PSTP-Scan models on the independent test set (‘Mix’ dataset). (H) Performance of PSTP-Scan block with different window sizes. Unlike Fig. 3 D to G, where the default three parallel PSTP-Scan blocks were used, each model here represents a single-block PSTP-Scan with a specific window size. The ‘kernel’ model configuration refers to passing the averaged vector directly to the trained MLP kernel in the default PSTP-Scan setup to generate output during test set evaluation.

**Fig. S3.**
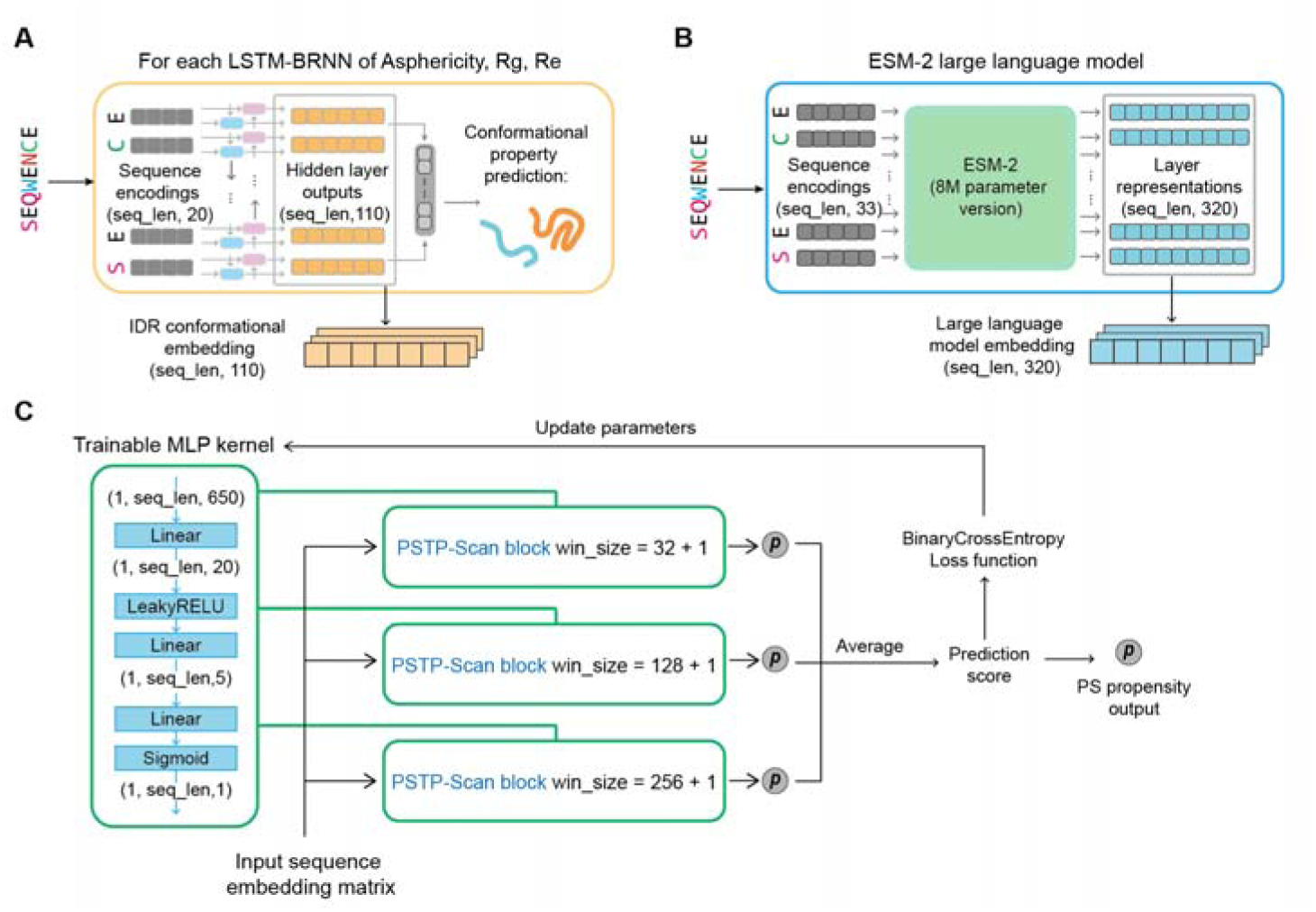
Schematic of the PSTP model, related to Fig. 1. (A) IDR conformational embedding: For each LSTM-BRNN sub-model in ALBATROSS, the sequence-length x 110-dimension hidden layer output is extracted prior to predicting the final property value. (B) Large language model embedding: Residue-level layer representations from the ESM-2 model (8 million parameters version) are extracted, resulting in a sequence-length x 320-dimensional matrix for each input sequence. (C) PSTP-Scan model training and inference: Three parallel PSTP-Scan blocks with identical architectures but different average pooling window sizes are employed to capture sequence information at varying local lengths. The predicted sequence-level propensities (single-value output from each block) are averaged to produce the final output. This output is optimized via backpropagation to update the weights of the MLP.

**Fig. S4.**
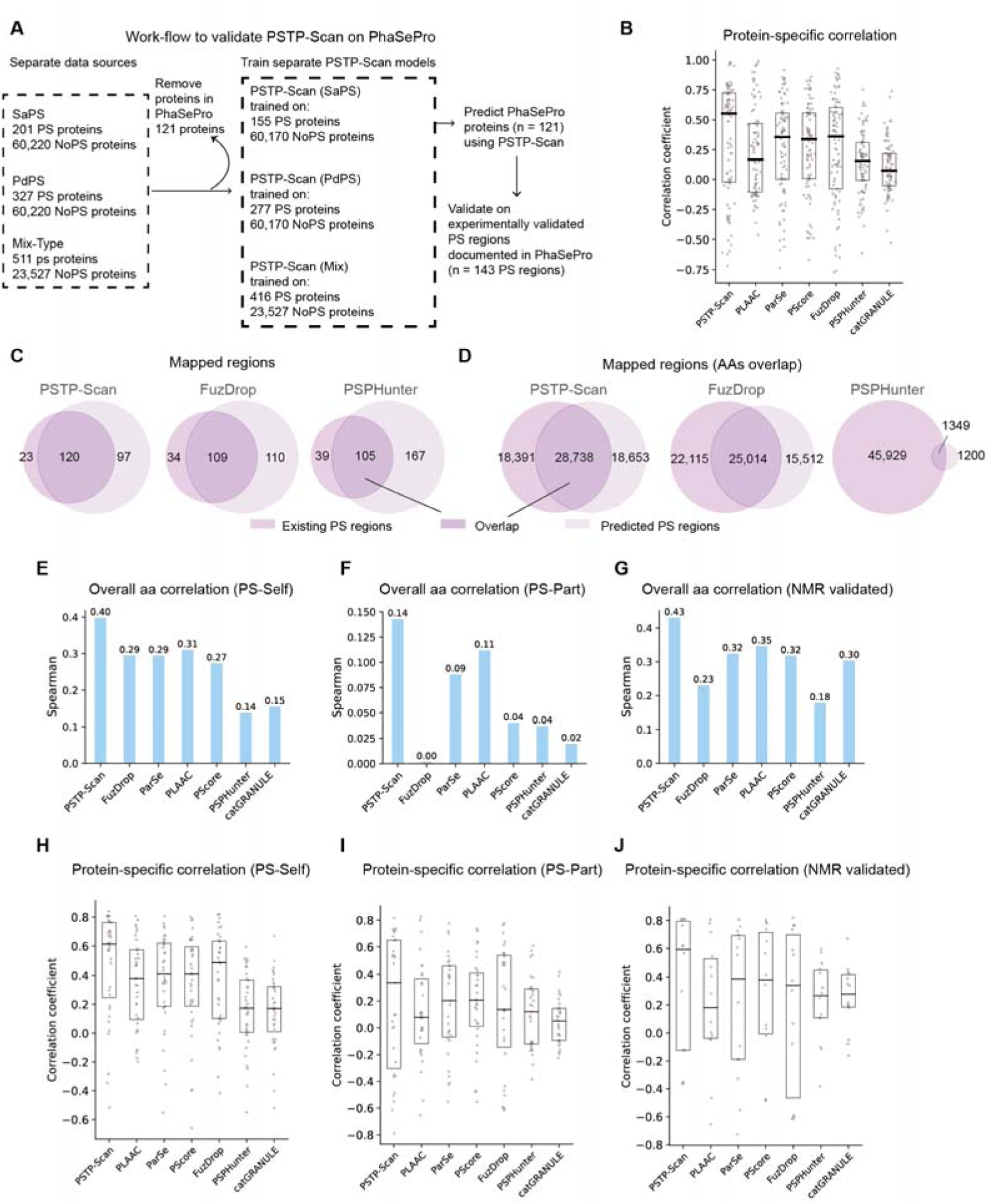
Evaluation of residue-level prediction scores for proteins documented in the PhaSePro database, related to Fig. 4. (A) Illustration of the validation process for PSTP-Scan in predicting PS driving regions. In this test, Each PSTP-Scan model was specifically trained excluding proteins documented in PhaSePro to prevent self-validation and data leakage. (B) Distribution of the protein-level Spearman correlation coefficients computed between the residue-level scores and the binary vector representing the PS region(s) for each protein, corresponding to Fig. 4E. Scatter points represent individual coefficient values, while the boxplot components within each violin, from top to bottom, represent upper quartile, median, lower quartile. (C and D) Comparison of PS regions predicted by PSTP-Scan, FuzDrop, and PSPHunter against those documented in PhaSePro, related to Fig. 4B. FuzDrop was trained using PhaSePro proteins. This panel illustrates the overlap between predicted regions (C) and the corresponding overlap at the individual amino acid level (D). (E and H) Parallel evaluations to Fig. 4C (E) and Fig. S4B (H), focusing exclusively on PS regions of PS-Self proteins (58 records). (F and I) Parallel evaluations to Fig. 4C (F) and Fig. S4B (I), focusing exclusively on PS regions of PS-Part proteins (50 records). (G and J) Parallel evaluations to Fig. 4C (G) and Fig. S4B (J), focusing exclusively on NMR-validated PS regions (14 records).

**Fig. S5.**
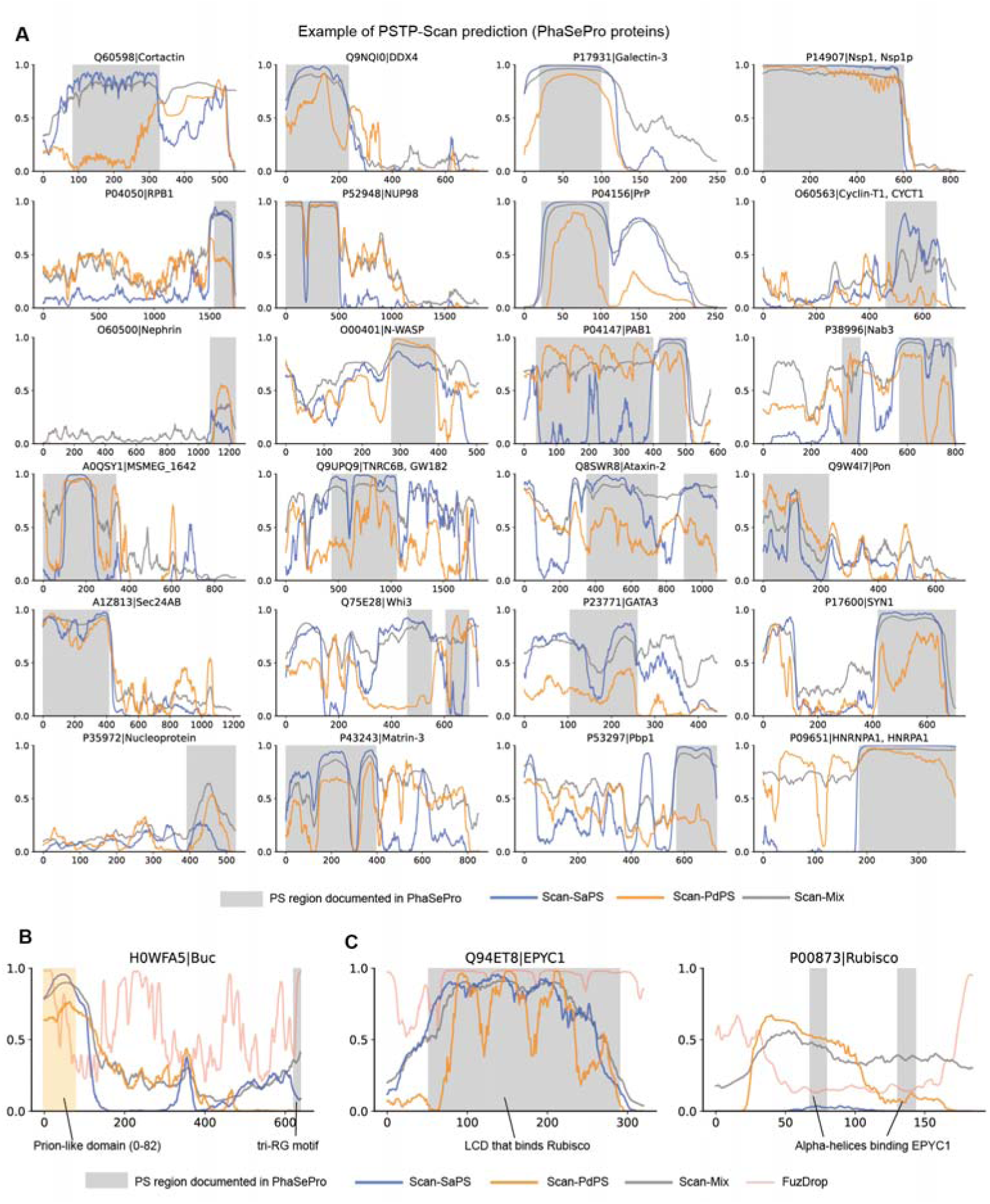
Examples of residue-level scores predicted by three PSTP-Scan models, related to Fig. 4, with experimentally validated PS regions highlighted in gray. RRM: RNA-recognition motifs.

**Fig. S6.**
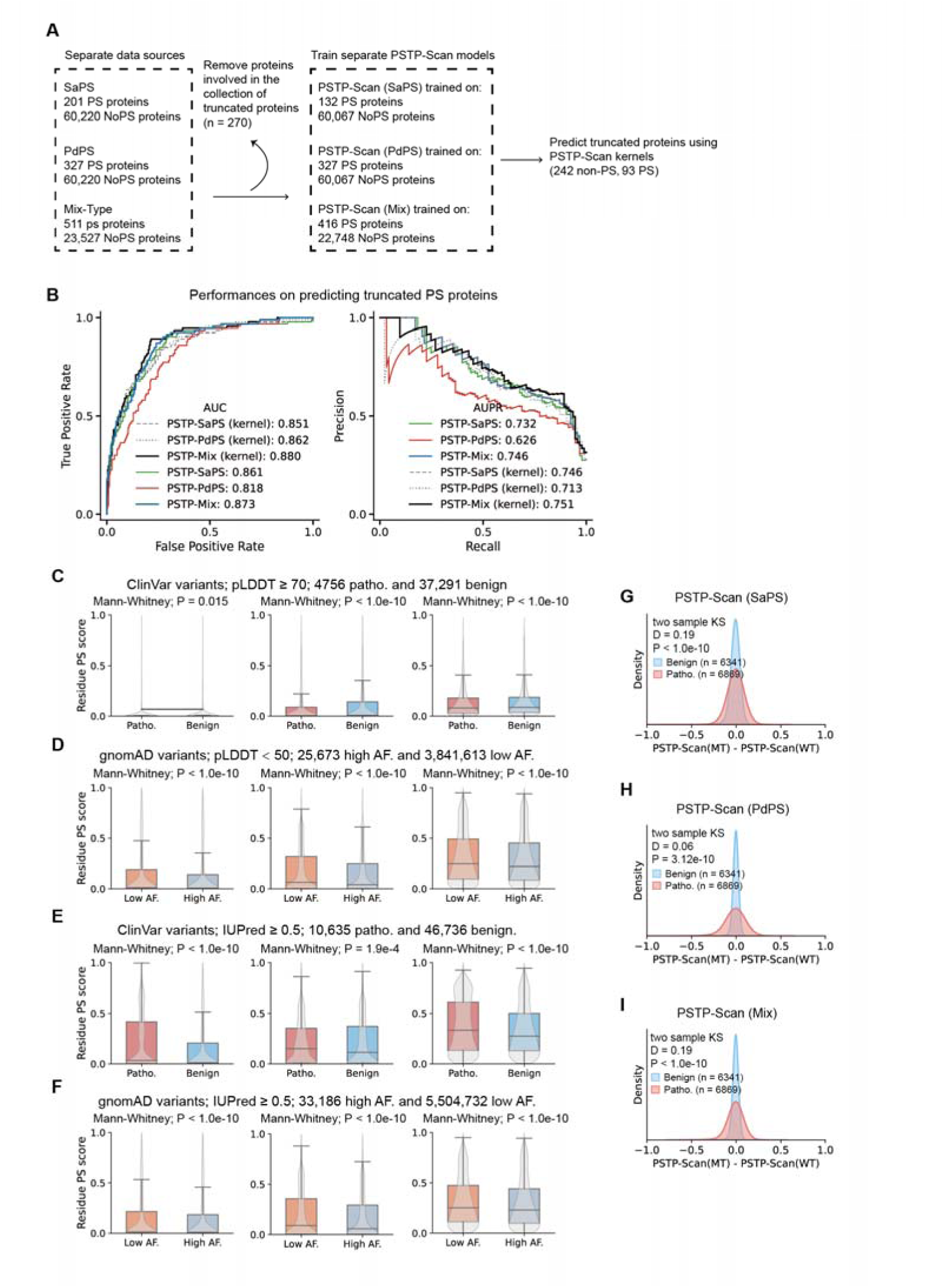
Extended scope of PS protein prediction and analysis of disease variants, related to Fig. 6. (A) Illustration of the validation process for PSTP-Scan in predicting truncated proteins. In this test, PSTP-Scan models were specifically trained after excluding proteins that overlap (share the same UniProt entry) with those in the truncated protein validation dataset. (B) AUC and AUPR evaluations of different PSTP-scan models for predicting truncated PS proteins. ‘kernel’ refers to using only the trained MLP kernel from the corresponding PSTP-Scan model to predict based on the averaged feature vector of each protein. PSTP-SaPS, PSTP-PdPS, and PSTP-Mix refer to the three trained PSTP-Scan models, respectively, with a single block applied and the window size set to 32+1. (C) PSTP residue-level scores for high pLDDT (pLDDT ≥ 70) pathogenicity and benign ClinVar variants. (D) PSTP residue-level scores for variants with high allele frequency (AF) and low allele frequency (AF) from gnomAD, located within low pLDDT (pLDDT < 50) regions. (E) PSTP residue-level scores for pathogenicity and benign ClinVar variants located in disordered regions as defined by IUPred3 (IUPred ≥ 0.5). (F) PSTP residue-level scores for high AF and low AF variants from gnomAD, located in disordered regions as defined by IUPred3 (IUPred ≥ 0.5). (H-I) A parallel evaluation to Fig. 6E but focusing on in-frame deletion. Pathogenic variants show significantly higher PSTP-Scan score variations compared to benign variants.

**Fig. S7.**
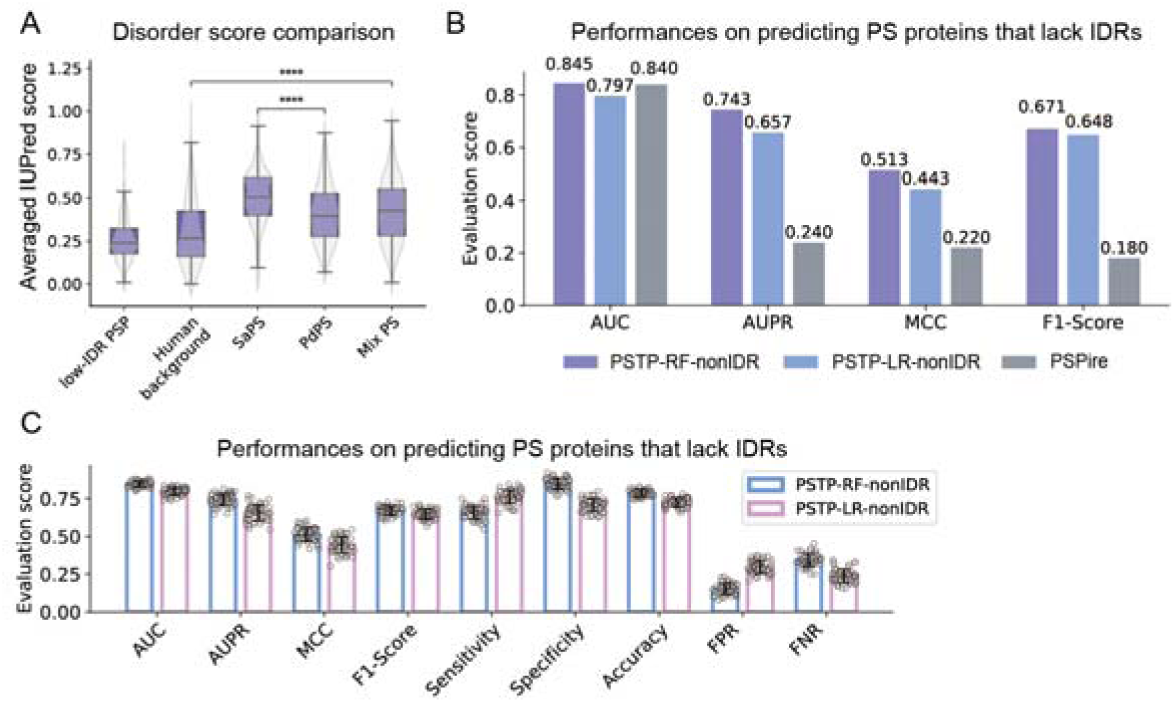
Model performance in predicting PS proteins lacking IDRs. (A) Distribution of IUPred3 scores, reflecting protein disorder across different types of proteins. PSP: PS protein; Low-IDR PSP: PS Proteins that lack IDRs; Human background: Proteins from the human proteome; SaPS: Self-assembly PS proteins; PdPS: Partner-dependent PS proteins; Mix PS: Mixture of different types of PS proteins. (B and C) Performance of PSTP-RF (PSTP-RF-nonIDR) and PSTP-LR (PSTP-LR-nonIDR) in predicting PS proteins. Both models were evaluated using the same training and independent datasets used for validating PSPire^69^. A head-to-head comparison was performed between PSTP models and PSPire, with the performance of PSPire extracted from the literature^69^. PSTP models were trained exclusively using the large language model features. For the computation of F1-Score, Precision, Matthews Correlation Coefficient (MCC), Accuracy, False Negative Rate (FNR), False Positive Rate (FPR), and Sensitivity, we set the cutoff at 0.4.

## Notes

### Competing Interest Statement

The authors have declared no competing interest.

### Summary of Updates

Figure 2, 5 and 6 revised

